# Engrailed transcription factors direct excitatory cerebellar neuron diversity and survival

**DOI:** 10.1101/2023.11.30.569445

**Authors:** Anjana Krishnamurthy, Andrew S. Lee, N. Sumru Bayin, Daniel N. Stephen, Olivia Nasef, Zhimin Lao, Alexandra L. Joyner

## Abstract

The excitatory neurons of the three cerebellar nuclei (eCN) form the primary output for the cerebellar circuit. The medial eCN (eCNm) were recently divided into molecularly defined subdomains in the adult, however how they are established during development is not known. We define molecular subdomains of the eCNm using scRNA-seq and spatial expression analysis and show they evolve during embryogenesis to resemble the adult. Furthermore, the eCNm is transcriptionally divergent from the rest of the eCN by E14.5. We previously showed that loss of the homeobox genes *En1* and *En2* leads to death of a subset of embryonic eCNm. We demonstrate that mutation of *En1/2* in embryonic eCNm results in cell death of specific posterior eCNm molecular subdomains and loss of TBR2 (EOMES) expression in an anterior subdomain, as well as reduced synaptic gene expression. We further reveal a similar function for EN1/2 in mediating TBR2 expression, neuron differentiation and survival in the two other cerebellar excitatory neuron types. Thus, our work defines embryonic eCNm molecular diversity and reveals conserved roles for EN1/2 in the cerebellar excitatory neuron lineage.

## INTRODUCTION

Critical to defining the development and function of an organ is determining the molecular heterogeneity of cell types and identifying the cellular properties of each subtype. The cerebellum provides an excellent system for exploring this question as it has only 3 excitatory cell types in addition to several inhibitory neurons, and the two lineages are derived from distinct embryonic progenitor zones (Leto et al., 2016; Joyner and Bayin, 2022). The adult cerebellum has an outer folded cerebellar cortex consisting of lobules situated above three bilaterally symmetrical groups of neurons called the cerebellar nuclei (CN) (Glickstein and Voogd, 1995; Voogd and Glickstein, 1998). The two groups of CN are subdivided into medial, intermediate and lateral nuclei (CNm, CNi and CNl) based on their location (Kebschull et al., 2023; Paxinos and Watson, 2006). There is preservation of a spatial map between the cells in the cortex and the CN, such that inhibitory Purkinje cells in the cortex located from medial to lateral project their axons to corresponding medial to lateral CN neurons (Apps et al., 2018). The CN are composed of excitatory cerebellar nuclei neurons (referred to as eCN) that have long-range projections to the rest of the brain/spinal cord, as well as inhibitory neurons (IN) that primarily project locally (Fujita et al., 2020; Judd et al., 2021; Kebschull et al., 2020; Kebschull et al., 2023). The CN are thus the primary output neurons of the cerebellum and mediate its wide array of motor and non-motor functions. During development, the eCN play a pivotal role in ensuring the proper expansion of several cell types in the lobules and thus growth of each lobule is dependent on accurate mapping of Purkinje cells to specific eCN (Willett et al., 2019). Given that the position of eCN in 3-dimensional (3D) space relates to the final morphology of the cerebellar cortex, it is critical to determine whether there is an underlying molecular map of the developing eCN that provides a framework for growth and circuit formation in the lobules.

The eCN are the first-born neurons of the cerebellum, arising during embryonic day (E)9.5-12.5 from an excitatory neuron progenitor zone called the rhombic lip (RL), soon after the cerebellar anlage is specified (Machold and Fishell, 2005; Wang et al., 2005). The newborn eCN migrate tangentially along the surface of the cerebellum and reside in the anterior-dorsal portion of the cerebellum in a region called the nuclear transitory zone (NTZ) until E15.5 (Altman and Bayer, 1985). They then rearrange to form the bilaterally symmetrical sets of 3 three nuclei. Importantly, eCN promote survival of their upstream presynaptic partners Purkinje cells (Willett et al., 2019). The Purkinje cells then stimulate proliferation of postnatal progenitors of the excitatory granule cells, interneurons and astrocytes of the cerebellar cortex through secretion of Sonic hedgehog (Corrales et al., 2006, Fleming et al., 2013). The eCN together with the Purkinje cells thereby regulate the size of the cerebellum (Willett et al., 2019).

Each nucleus of the adult mouse CN has molecularly defined subdomains based on their transcriptome and location. Using single nuclear RNA sequencing (snRNA-seq) of dissected mouse CN, it was shown that eCN in all three nuclei (eCNm, eCNi and eCNl) can be divided into two classes of neurons differing in size, electrophysiological properties and gene expression profiles, and further subdivided into spatially distinct subpopulations within each nucleus (Kebschull et al., 2020). Curiously, they found that the eCNm are especially distinct from the eCNi and eCNl that are transcriptionally more similar to each other. In a separate study, the mouse adult eCNm was subdivided into 4 subpopulations based on the expression of 3 marker proteins (SPP1, SNCA and CALB2) and their distinct location and neural circuit organization (Fujita et al, 2020) (Fig. 1A). Similar to the snRNA-seq analysis, the four subpopulations were found to have different cell sizes and spatial distributions within the eCNm. Although these studies provide insights into the molecularly defined subpopulations of the adult eCN, how and when this molecular diversity emerges during development remains to be determined.

**Figure 1:**
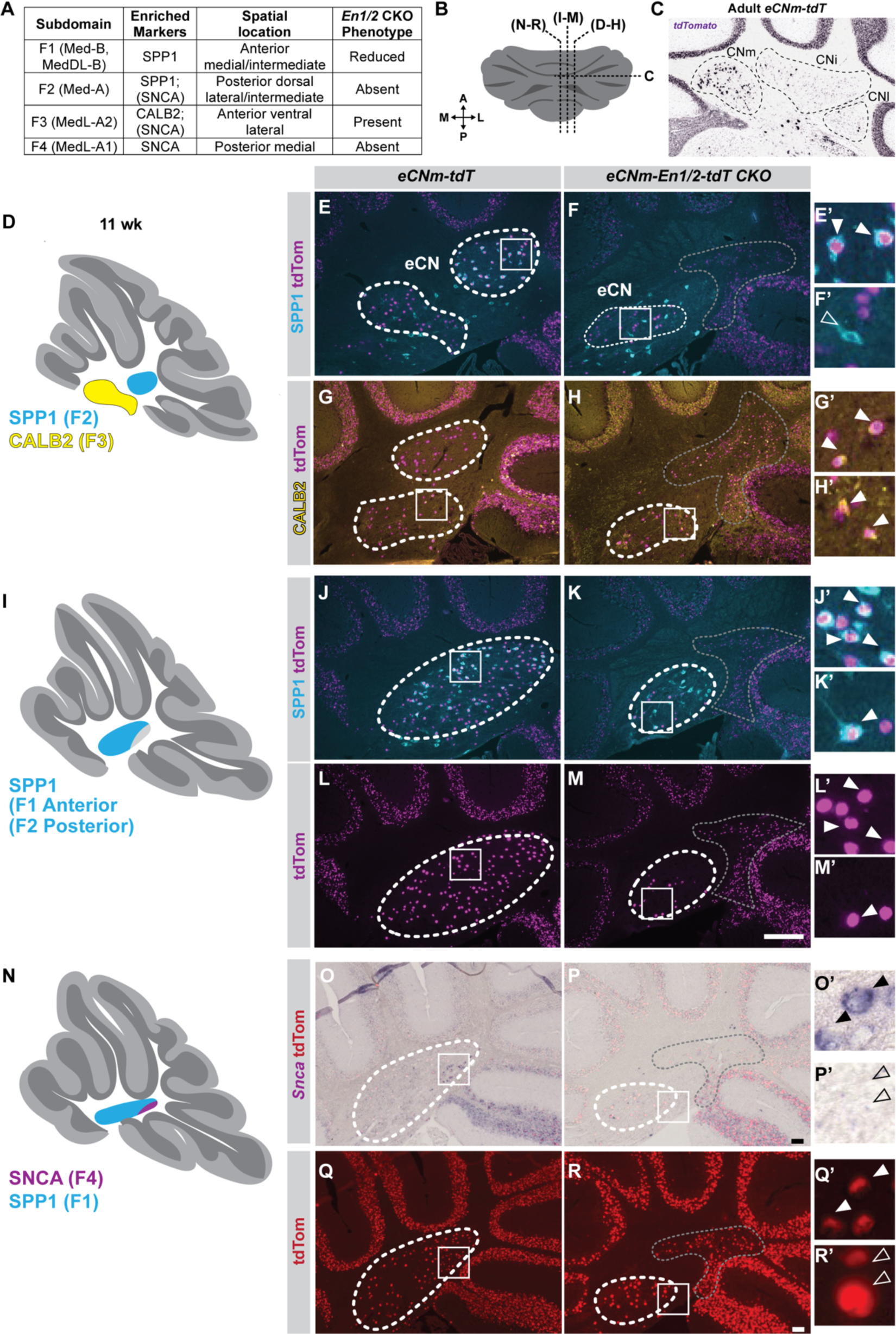
Two posterior subpopulations of the eCNm, defined by SPP1 laterally and SNCA medially, are absent in *eCNm-En1/2-tdT* CKO animals and anterior domains are diminished. (A) Table listing adult eCNm populations defined by Fujita et al, 2022 and Kebschull et al., 2020. (B) Cerebellar schematic indicating levels of (C-R). (C) ISH for tdT in Allen Brain Mouse Connectivity Atlas of SepW1-Cre_NP39:Ai14 experiment 488246145 anterior coronal section with CN outlined. (D-R’) IF analysis of SPP1 and CALB2 and RNA in situ analysis of *Snca* as indicated in the eCNm on sagittal sections at lateral levels of the cerebellum of 11 week *eCNm-En1/2-tdT* CKOs and controls. Squares indicate location from which higher power images (indicated by prime symbol) were taken. Solid arrowheads: tdT+ SPP1+ cells in (E’), tdT+ CALB2+ cells in (G’, H’), tdT+ SPP+ cells in (J’-M’). Empty arrowheads: tdT-SPP1+ cells in (F’), tdT+ *Snca-* cells in (P’,R’). White dashed outlines: eCNm, grey dashed outlines: ectopically positioned UBCs. Scale bars: 100 μm.

The homeobox genes Engrailed1/2 (*En1/2*) encode transcription factors that are key regulators of mouse cerebellar growth, patterning, and circuit formation (Cheng et al., 2010; Joyner, 1996; Joyner et al., 1991; Kuemerle et al., 1997; Millen et al., 1994; Millen et al., 1995; Orvis et al., 2012; Sgaier et al., 2007; Sillitoe et al., 2008; Sillitoe et al., 2010; Willett et al., 2019; Wurst et al., 1994), as well of survival of specific mid- and hindbrain neuron types (Altieri et al., 2015; Fox and Deneris, 2012; Sgadò et al., 2006; Simon et al., 2001). These genes therefore provide a powerful entry point to study cerebellar development. Of interest for dissecting molecular characteristics of the eCN, we showed that ablation of *En1/2* in eCN leads to death of a subset of eCNm and eCNi between E15.5 and 17.5 with a preferential loss of posteriorly located neurons (Willett et al., 2019). Thus, one function of EN1/2 is to promote the survival of specific subpopulations of eCN. This result also indicates there could be molecular diversity of the eCN as early as E15.5. However, whether embryonic eCN subdomains can be defined and how they evolve and are related to the subpopulations of eCN defined in the adult is not known.

Given the recently defined subpopulations of the mouse adult eCNm and role of *En1/2* in regulating survival of some posterior eCNm, we focused our current study on defining the molecular diversity of the embryonic eCNm and its temporal evolution in normal embryos and those lacking *En1/2*. Using single cell RNA-sequencing (scRNA-seq) and immunohistochemical spatial analysis, we reveal that the eCNm is transcriptionally distinct from the eCNi+eCNl by E14.5 (shortly after eCN neurogenesis). Furthermore, the spatial expression domains of six proteins specific to the eCNm (TBR1, CALB2, EOMES/TBR2, PAX6, BARHL1 and LMX1A) become refined during embryonic development and at E17.5 their combined expression defines 4 subdomains similar to those in the adult. scRNA-seq analysis of E14.5 mutants lacking *En1/2* in all eCN prior to their cell death revealed EN1/2 promote expression of synaptic genes in eCNm. *En1/2* also regulate patterning of an anterior TBR2-expressing eCNm region, but cell viability of two posterior BARHL1+ eCNm subdomains. Uncovering a general role for EN1/2, the proteins upregulate TBR2 expression in the two other excitatory derivatives of the rhombic lip (GCPs and unipolar brush cells (UBCs)) and promote UBC viability and differentiation of UBCs and GCPs. Our study thus defines 4 spatially segregated eCNm subpopulations in the embryo and demonstrates EN1/2 play sequential roles in first setting up spatial patterning of the eCNm and later promoting neural specification and/or viability of specific subpopulations of all excitatory cerebellar neurons.

## RESULTS

### Adult eCNm subdomains are differentially vulnerable to embryonic loss of *En1/2*

As a step towards gaining a deeper understanding of the molecular subdivisions of the adult eCN, we determined whether the neurons lost in the eCNm of *En1/2* conditional mutants (CKOs) belong to one of the four spatial subdomains that can be identified based on preferential expression of SPP1, CALB2 and/or SNCA expression (F1-4; Fujita et al., 2020) (Fig.1A). We performed immunofluorescence (IF) and in situ hybridization (ISH) analysis on cerebellar sagittal sections of adult CKOs lacking *En1/2* in all eCN and the GCPs and that carry a nuclear tdTomato (tdT) reporter (*Atoh1-Cre/+; En1^lox/lox^, En2^lox/lox^*; *R26^LSL-tdT/+^* mice), referred to as *eCN+GCP-En1/2-tdT* CKOs, and littermate controls (*En1^lox/lox^; En2^lox/lox^*; *R26^LSL-tdT/+^*), referred to as *En1/2^f/f^* mice (Fig. S1). We also analyzed a new allele in which *En1/*2 are deleted specifically in the eCNm using a Selenow *Bac* construct to drive *Cre* (Gerfen et al., 2013) (*SepW1-Cre/*+; *En1^lox/lox^, En2^lox/lox^*; *R26^LSL-tdT/+^* mice or *eCNm-En1/2-td* CKOs) and controls in which the eCNm were labeled using *SepW1-Cre/*+ with tdT *(eCNm-tdT*) (n=3 animals per genotype; Fig. 1B-R’). Additional details of the *SepW1-Cre* lineage and adult *En1/2* mutant phenotype will be described elsewhere (Lee at al. in preparation). Strikingly, at all mediolateral levels, the posterior domain of the eCNm was absent. At lateral and intermediate levels of *eCNm-En1/2-tdT* CKO cerebella, where SPP1 alone marks most of the cells in the posterior dorsal region (F2) of the eCNm, no SPP1+/tdT+ cells were detected, whereas in *eCNm-tdT* controls a large distinct tdT+ domain was present (Fig. 1D-F’). The anterior and ventral region of the eCNm, which contains SPP1+ cells at medial and intermediate levels (F1) and scattered CALB2+ neurons more laterally (F3) was reduced in size in the *eCNm-En1/2-tdT* CKO mutants compared to *eCNm-tdT* controls (Fig. 1G-M’). At the most medial level, where *Snca* normally labels the posterior dorsal region of the eCNm (F4), no *Snca* mRNA was detected in *eCNm-En1/2-tdT* CKO mice and the tdT+ posterior dorsal eCNm domain was absent (Fig. 1N-R’). Similar results were seen in *eCN+GCP-En1/2-tdT* CKOs and littermate controls (Fig. S1). Thus, absence of *En1/2* in eCN results in loss of the SPP1+ and *Snca* posterior dorsal domains (F2 and F4) of the eCNm throughout the mediolateral axis and a partial loss of the remaining SPP1+ or CALB2+ anterior ventral eCNm (F1 and F3).

### In the embryonic NTZ TBR1 and OLIG2 mark complementary medial and lateral eCN domains, respectively

As a basis for determining whether the subpopulations of eCNm that die in *En1/2* eCN conditional mutants are transcriptionally distinct embryonically, we defined the transcriptional landscape of control eCN at E14.5, when they still reside in the NTZ, and at E17.5, when the three nuclei can be distinguished. We performed scRNA-seq on tdT+ cells isolated by fluorescence activated cell sorting (FACS) from E14.5 *eCN+GCP*-*tdT* embryos (n=2 pools) using a 10X Genomics platform (Fig. S2A, B). After filtering out of low-quality cells and integrating both replicates using Seurat v4.1 (see Methods), a total of 9,620 cells were obtained (Fig. 2A, Fig. S2C). Unsupervised clustering of the combined cells yielded 10 clusters that as expected were either enriched for *Meis2* that marks the eCN (adjusted p-value ≤ 0.05, Table S1) or *Pax6* that is enriched in GCPs compared to eCN (adjusted p-value ≤ 0.05; Table S1)(Fig. 2A’-A”, Fig. S2D-F)(Willett et al, 2019; Morales and Hatten, 2006; Engelkamp et al., 1999a). Outside the cerebellum, *Meis2* is expressed in neurons of the anterior NTZ (Fig. S2G, H) destined for the mid/hindbrain, including the isthmic nuclei (Wizemann et al, 2019), as well as in cell populations adjacent to the cerebellum (Fig. S2G-I). To identify which of the *Meis2-* enriched clusters were eCN, we isolated clusters 2, 3, 4 6 and 7 and reiterated unsupervised clustering (Fig. 2B). 9 clusters were obtained, of which the largest cluster (0) was marked by *Tbr1* (808 cells) and two other clusters (1 and 7) were enriched for *Olig2* (1,040 cells total) (Fig. 2B-B”). *Tbr1* and *Olig2* were reported to be expressed in the E13.5 NTZ (Wizemann et al, 2019) and at E18.5 to mark the eCNm and eCNi + eCNl, respectively (Ju et al, 2016, Seto et al, 2014). The remaining clusters appear to contain cells outside the cerebellum based on the expression patterns of their significantly enriched markers in the Allen Developing Mouse Brain Atlas E13.5 and 15.5 RNA ISH datasets (Table S2, Fig. S2G-I). We therefore excluded clusters 2-6 and 8, and repeated clustering on the *Tbr1+* and *Olig2+* clusters, which again yielded three clusters – one *Tbr1* expressing cluster (0) and two *Olig2+* clusters (Fig. 2C-C”) which were enriched for either *Nrp1* (1) or *Nr2f1* (2) (Fig S2J).

**Figure 2:**
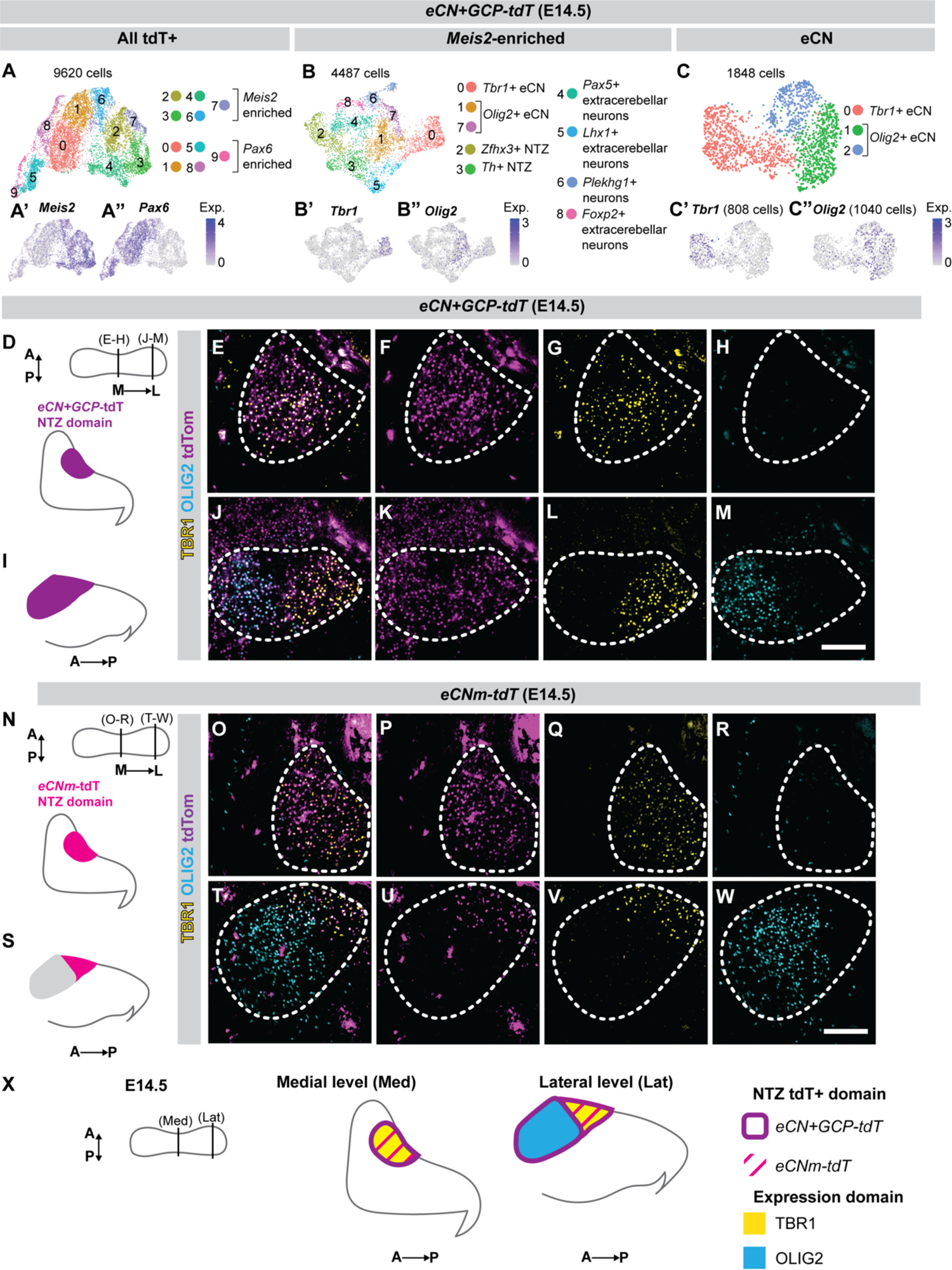
Error! No text of specified style in document.: TBR1 and OLIG2 label complementary medial and lateral domains, respectively within the eCNm containing region of the NTZ. (A) Uniform Manifold Approximation and Projection (UMAP) visualization of tdT+ cell clusters from *eCN+GCP-tdT* E14.5 cerebella (n = 2 pools), with *Meis2* (A’) and *Pax6* (A”) expression. (B) UMAP visualization of *Meis2-*enriched cluster re-clustering, with *Tbr1* (B’) and *Olig2* (B”) expression. (C) UMAP visualization of *Tbr1+* and *Olig2+* reclustering with *Tbr1* (C’) and *Olig2* (C”) expression. (D) Schematic of an E14.5 cerebellum showing levels of (E-H) and (J-M) sections and below a medial sagittal section. Merge (E) and tdT (F), TBR1 (G) and OLIG2 (H) single channel IF images of medial E14.5 *eCN+GCP-tdT* NTZ. (I) Schematic of a lateral sagittal section. Merge (J) and tdT (K), TBR1 (L) and OLIG2 (M) single channel IF images of lateral E14.5 *eCN+GCP-tdT* NTZ. (N) Schematics of E14.5 cerebellum showing levels of (O-R) and (T-W), and below a medial sagittal section. Merge (O) and tdT (P), TBR1 (Q) and OLIG2 (R) single channel IF images of medial E14.5 *eCNm-tdT* NTZ. (S) Schematic of a lateral sagittal section. Merge (T) and tdT (U), TBR1 (V) and OLIG2 (W) single channel IF images of lateral E14.5 *eCN+GCP-tdT* NTZ. Dashed outlines: eCN containing domain of NTZ. (X) Schematic summary of expression results. Scale bars: 100 μm (E-M), (O-W).

Having identified E14.5 eCN clusters, we confirmed that TBR1 and OLIG2 were indeed expressed in *eCN+GCP-Cre* derived cells by performing IF on cerebellar sagittal sections across the mediolateral extent of E14.5 *eCN+GCP-tdT* embryos. TBR1 and OLIG2 were detected in complementary domains throughout the tdT+ NTZ (Fig. 2D-M). TBR1 was present in the entire NTZ at medial levels (Fig. 2E, G), and was restricted to the posterior NTZ laterally (Fig. 2J, L). In contrast, OLIG2 was absent from the medial NTZ (Fig. 2E, H) and marked the anterior NTZ, complementary to TBR1, at lateral levels (Fig. 2J, M). Since we observed some TBR1+ eCN at lateral levels, we tested whether TBR1 is exclusive to the eCNm at this stage by analyzing E14.5 *eCNm-tdT* cerebella (Fig. 2N-W). Strikingly, we found that TBR1 was co-expressed with nearly all tdT+ cells in *eCNm-tdT* cerebella (Fig. 2O,Q,T,V), whereas OLIG2 was excluded from tdT+ cells (Fig. 2O,R,T,W). As the *eCNm-Cre* transgene marks only the eCNm in adult mice, these results demonstrate that eCNm cells are transcriptionally distinct from the remaining eCNi+eCNl shortly after they are generated and are uniquely marked by TBR1. Furthermore, some eCNm neurons are initially located laterally (Fig. 2X).

### Markers enriched in E14.5 eCNm have distinct anteroposterior and mediolateral distributions

We next generated a list of genes enriched in the *Tbr1+* eCNm cluster 0 at E14.5 (Table S3). Interestingly, of the three genes used to subdivide the adult eCNm into subpopulations, only *Calb2* was enriched in the *Tbr1+ eCNm* cluster in our scRNA-seq dataset (Fig. 3A). *Snca* was present in a few cells in both the eCNm and eCNi + eCNl clusters (Fig. S3A) and the Allen Developing Mouse Brain Atlas shows *Snca* broadly in the eCN at E15.5 and E18.5 (Fig. S3B-L). *Spp1* was not detected in our dataset or using IF analysis of E14.5 or E17.5 *eCNm-tdT* cerebellar sections (Fig. S3M-V). Thus, our expression results indicate that the transcriptome of the embryonic eCNm is distinct from that of the mature eCNm and thus must evolve during postnatal development.

**Figure 3:**
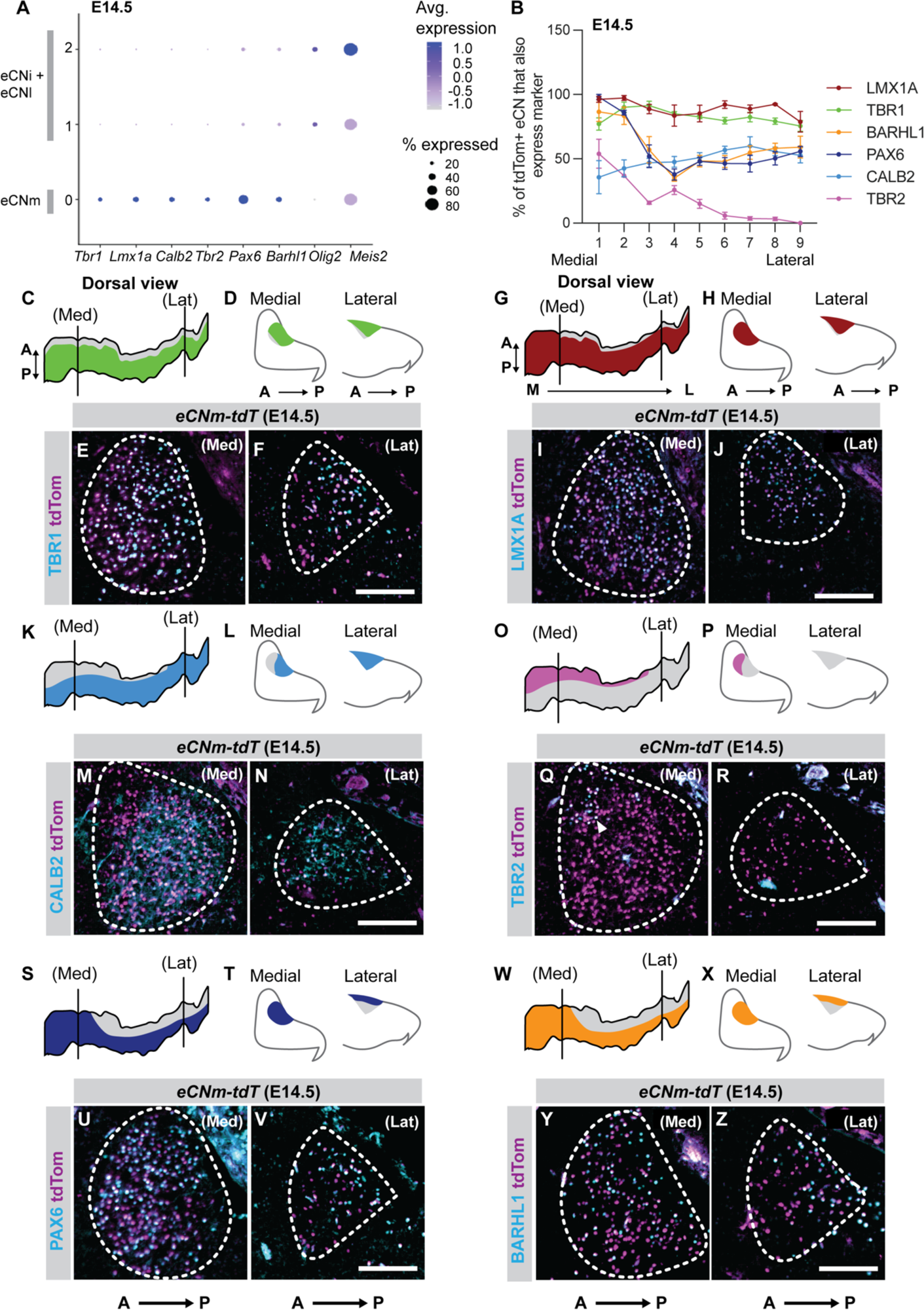
Protein markers enriched in eCNm have distinct spatial patterns in the NTZ at E14.5. (A) Expression of eCNm, eCNi+eCNl and pan-eCN markers across all eCN clusters. (B) Percentage of double positive (tdT and eCNm marker) cells in the NTZ at nine medio-lateral levels in the E14.5 *eCNm-tdT* cerebellum (n = 3 animals). Data is presented as mean ± s.e.m. (C-Z) Colocalization of tdT and each eCNm marker in E14.5 *eCNm-tdT* cerebella at medial and lateral levels, with schematics of marker expression projected onto a dorsal view of E14.5 eCNm 3D-reconstruction (C, G, K, O, S, W) and sagittal section schematics (D, H, L, T, P, X). Dashed outlines: eCNm. Scale bars: 100 μm.

To determine if molecular subpopulations of the eCNm exist at E14.5, we selected four significantly enriched eCNm marker genes (*Tbr2, Pax6, Barhl1* and *Lmx1a*; Table S3) in addition to *Calb2* and *Tbr1,* for spatial expression analysis because validated antibodies were available and they had been described as expressed in the NTZ or the eCNm (Chizhikov et al., 2006; Fink et al., 2006; Rose et al., 2009; Yeung et al., 2016) (Fig. 3A). We quantified the percentage of tdT+ eCNm NTZ cells in *eCNm-tdT* E14.5 embryos positive for each marker on every tenth sagittal section from medial to lateral levels (Fig. 3B). To represent the expression domain of each marker in the anteroposterior plane, we performed a 3D reconstruction of the E14.5 eCNm-tdT+ cells and projected the expression of each marker onto a dorsal view of the reconstruction. TBR1 (Fig. 3C-F) and LMX1A (Fig. 3G-J) were expressed across most of the tdT+ eCNm except for a few anteriorly located cells. Strikingly, the other markers were expressed in distinct domains restricted along the mediolateral and/or anteroposterior axes. We found that CALB2 (Fig. 3K-N) labels a subset of cells across the mediolateral extent of the tdT+ NTZ, aside from some anterior cells at medial levels where TBR2 (Fig. 3O-R) is expressed. PAX6 (Fig. 3S-V) and BARHL1 (Fig. 3W-Z) were enriched in the posterior-most region of the tdT+ NTZ laterally and present in most tdT+ cells medially. Thus, we demonstrate that the eCNm already have molecular subpopulations at E14.5 that correlate with their 3D position in the NTZ, suggesting subpopulations of neurons are transcriptionally defined soon after they are generated.

### eCNm molecular subpopulations evolve between E14.5 and E17.5

We next asked whether the molecular diversity of the eCNm at E14.5 is stable or evolves during development and establishment of the eCN nuclei. We generated 3D reconstructions of the expression domains of the same set of markers from sagittal cerebellar sections at E17.5, the earliest stage when CN morphology and positioning closely resembles that of the adult (Fig. S4). Interestingly, all six markers labeled a lower percentage of the tdT+ cells in *eCNm-tdT* E17.5 cerebella than at E14.5 (Fig. 4A), and the expression domains of most of the markers changed between the time points (compare Fig. 3B to Fig. 4B). Additionally, it became apparent that the expression domains of most markers developed additional spatial segregation along the dorsoventral axis. TBR1 was found to be slightly more enriched laterally than medially at E17.5, and throughout the mediolateral axis it appeared more highly expressed ventrally (Fig. 4C-F). The only marker that showed a very similar pattern of expression at E17.5 compared to E14.5 was TBR2 (Fig. 4G-J), which was expressed solely in a small group of anterior and ventral tdT+ eCNm, with enrichment at medial levels. At E17.5 CALB2 became enriched in the ventral regions of the tdT+ eCNm at lateral and medial levels (Fig. 4K-N). BARHL1 was highly enriched in the posterior dorsal eCNm at all levels, aside from an additional small group of anterior eCNm at medial levels (Fig. 4O-R). PAX6 (Fig. 4S-V) and LMX1A (Fig. 4W-Z) became restricted at E17.5 to anterior and ventral regions in the eCNm across the mediolateral axis, with the expression of PAX6 being more scattered than LMX1A. Thus, the expression domains of most embryonic eCNm markers evolve between E14.5 and E17.5. Based on combined expression patterns, the eCNm can be considered to have 4 subdomains at E17.5 related to those described in the adult (see Discussion).

**Figure 4:**
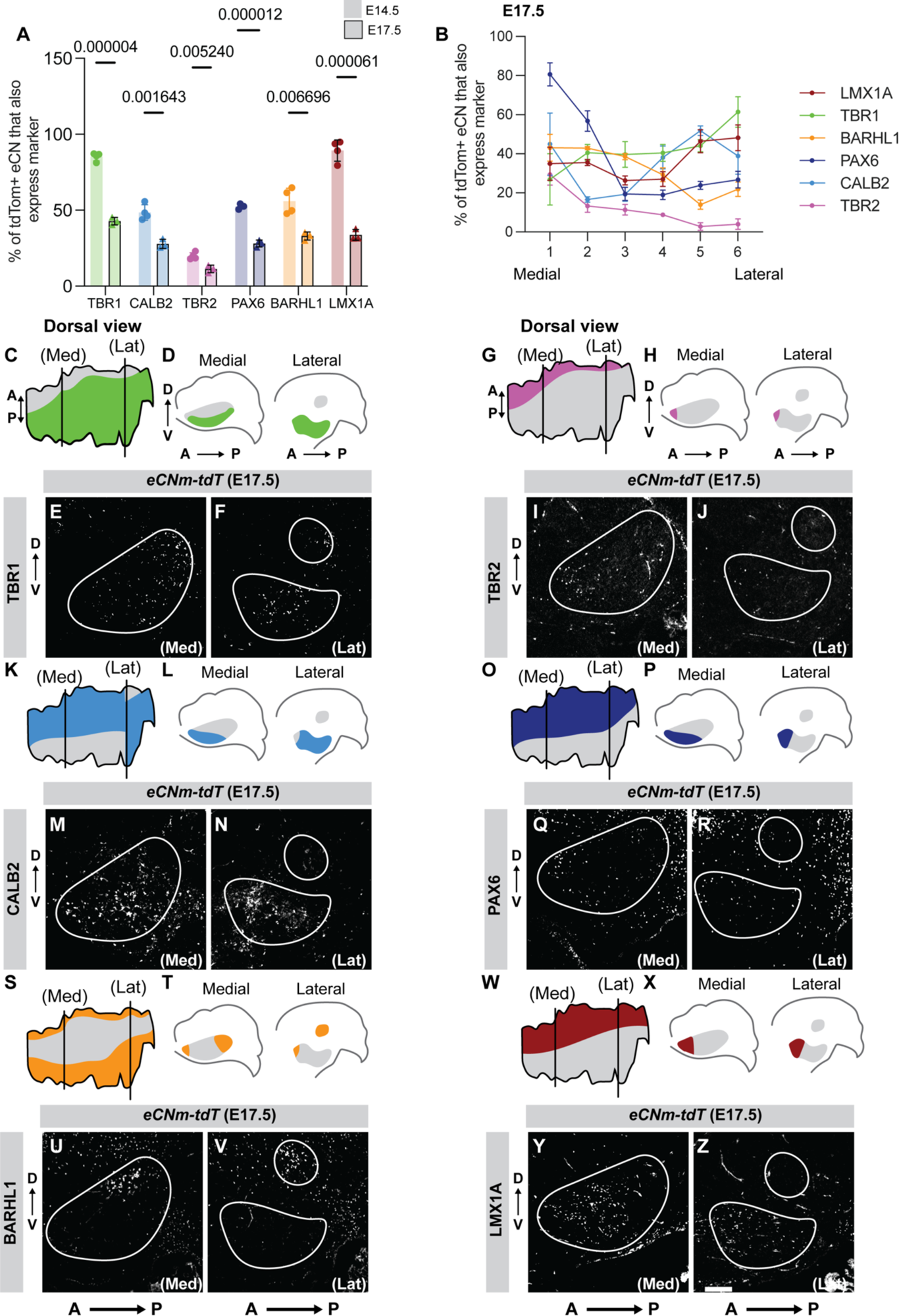
Protein markers enriched in eCNm evolve between E14.5 and E17.5. (A) Comparison of percentage of double positive (tdT and marker) eCNm at E14.5 and E17.5 in *eCNm-tdT* mice (n = 3 animals per age, Student’s t-test). (B) Percentage of double positive eCNm at six medio-lateral levels in the E17.5 *eCNm-tdT* cerebellum (n = 3 animals). Data is presented as mean ± s.e.m. (C-Z) Colocalization of tdT and each eCNm marker in E17.5 *eCNm-tdT* cerebella at medial and lateral levels, accompanied by schematics of eCNm marker expression projected onto a dorsal view of E17.5 eCNm 3D-reconstruction (C, G, K, O, S, W) and sagittal section schematics (D, H, L, T, P, X). Dashed outlines: eCNm. Squares in (C-Z) indicate location from which images in (C’- Z’) were taken. Scale bars: 100 μm.

### Initial segregation of eCNm and eCNi + eCNl is maintained in *En1/2* CKOs but neural maturation genes are reduced

Having established a spatial molecular map of the embryonic eCNm, we next asked whether loss of *En1/2* alters the initial patterning of the eCN into medial and eCNi+eCNl populations. scRNA-seq analysis on FACS isolated tdT+ cells from *eCN+GCP-En1/2-tdT* CKO samples (n=2 pooled samples with 5 embryos each) (Fig. S5). 8,606 cells passed quality control and formed 11 clusters (Fig. S5A-F, Table S4), of which *Meis2*-enriched cells (1925 cells) were further clustered (Fig. S5G-J, Table S5) to identify eCNm (805 cells) which contained three *Olig2*-enriched clusters and one *Tbr1* cluster (Fig. S5K-N, Table S6). The percentage of *Tbr1+* cells of all the *eCNm* in the mutant (*eCN+GCP-En1/2-tdT* CKO) dataset (48.14%) was similar to that of the control (*eCN+GCP-tdT)* dataset (43.72%) and like in controls the *Olig2*+ clusters in mutants were enriched for *Nrp1* or *Nr2f1* (Fig. S5N). To test whether the three *eCN+GCP-En1/2-tdT* CKO *Olig2+* clusters were similar to the two *eCN+GCP-tdT Olig2+* clusters (see Fig. 2C), we generated predicted identities for cells from the *eCN+GCP-En1/2-tdT* CKO clustering using the *eCN+GCP-tdT* dataset as the reference dataset. We found that most *Olig2+* cells in the *eCN+GCP-En1/2-tdT* CKO clustering mapped to one of the two *eCN+GCP-tdT Olig2+* clusters (Fig. S5O).

We then integrated our *eCN+GCP-En1/2-tdT* CKO samples with the *eCN+GCP-tdT* control samples (18,400 cells total) and performed reiterative unsupervised clustering to identify eCNm (Fig. 5A-I’, Fig. S6A-D, Table S7-9). The eCNm cells from the combine genotypes (2,780 cells) formed one *Olig2*+ and one *Tbr1*+ cluster and a small *Mafb*+ cluster that is a contaminant from outside the cerebellum (Fig. 5G-I’, Fig. S6E-F). The mutant and control cells were distributed similarly throughout the three clusters. The *Olig2+* cluster had two subdomains defined by markers specific to each of the two *Olig2+* control clusters (*Nrp1* and *Nr2f1*) indicating a further subdivision exists (Fig. S6G-O).

**Figure 5:**
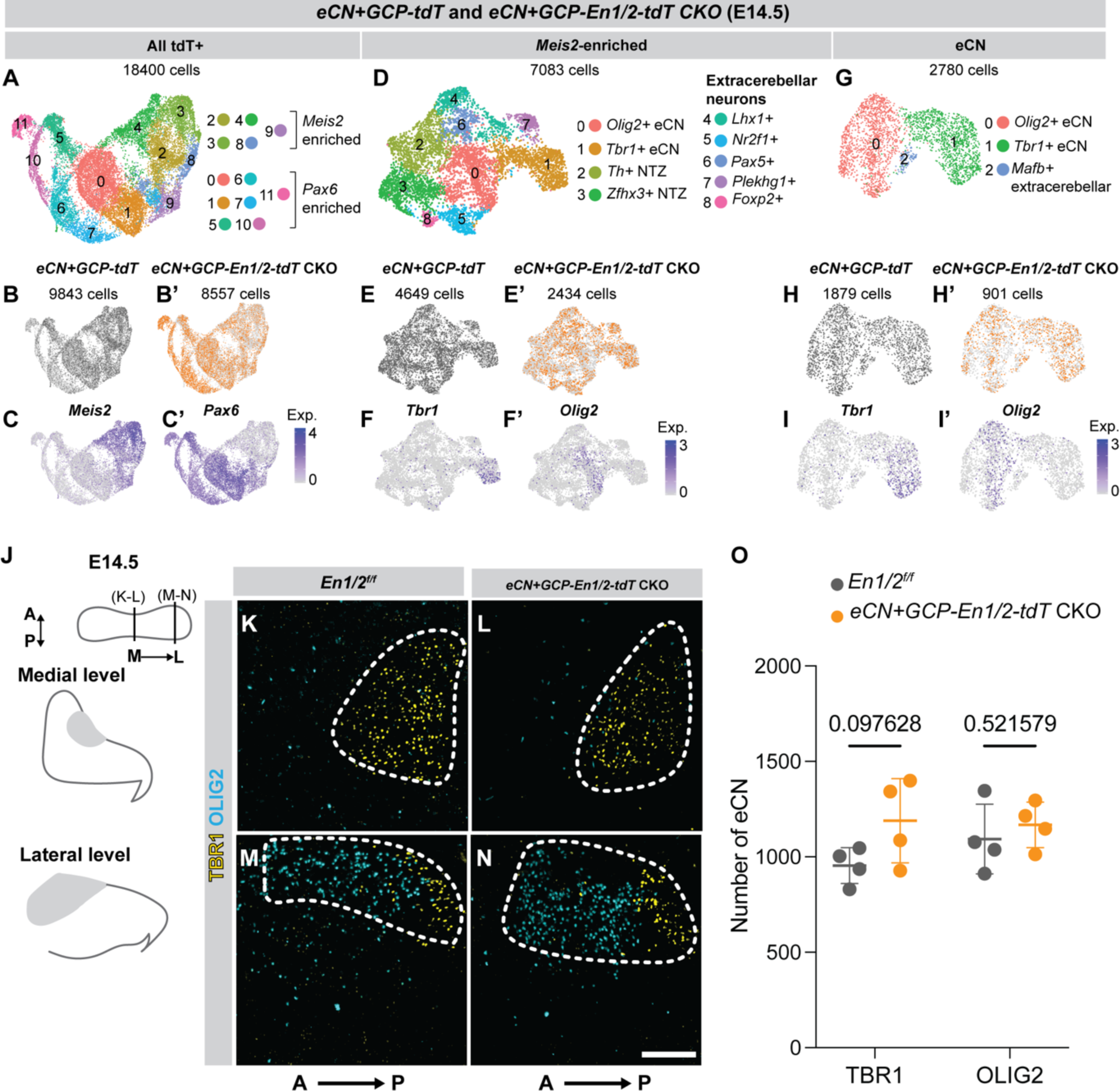
Initial patterning of eCN into complementary TBR1 and OLIG2 domains at E14.5 appears normal in *En1/2* CKOs. (A) UMAP visualization of tdT+ cell clustering of control and *eCN+GCP-En1/2-tdT* CKO eCN combined at E14.5. (B-B’) Distribution of cells from each genotype across clusters. (C-C’) Expression of *Meis2* and *Pax6.* (D) UMAP visualization of *Meis2-*enriched clusters reclustered. (E-E’) Distribution of cells from each genotype across clusters. (F-F’) Expression of *Tbr1* and *Olig2*. (G) UMAP visualization of reclustering of *Tbr1+* and *Olig2+* clusters. (H-H’) Distribution of cells from each genotype across clusters. (I-I’) Expression of *Tbr1* and *Olig2*. (J-N) Schematics and IF images of TBR1 and OLIG2 expression in the NTZ of E14.*5 En1/2^f/f^*(K,M) and *eCN+GCP-En1/2-tdT* CKO (L,N) sagittal sections of cerebella at medial and lateral levels of the cerebellum. Dashed outlines: eCN containing domain of NTZ. (O) Quantification of MEIS2+ TBR1+ or OLIG2+ cells on 9 sections across 3 NTZ (n = 4 animals per genotype; Student’s t-test). Scale bars: 100 μm (K-N).

IF staining on littermate control and *eCN+GCP-En1/2-tdT CKO* E14.5 cerebella (n = 4 animals per genotype) for TBR1 and OLIG2 across the CN revealed no obvious change in the mediolateral and anteroposterior distributions of the markers compared to controls (Fig. 5J-N). Additionally, quantification of TBR1+ and OLIG2+ eCN from every tenth sagittal section of the NTZ demonstrated no change in the number of cells expressing either marker between *eCN+GCP-En1/2-tdT* CKO and *eCN+GCP-tdT* embryos (Figure 5O). Thus, *En1/2* does not play a major role in the initial patterning of the eCN into eCNm and eCNi+eCNl populations.

Given there was no difference between the number or distribution of TBR1+ and OLIG2+ eCN, we then performed differential gene expression analyses between the *eCN+GCP-En1/2-tdT* CKO and *eCN+GCP-tdT* cells within the *Tbr1*+ or *Olig2*+ cluster using Libra, a pseudo-bulk differential gene expression algorithm (Squair et al., 2021)(Figure S7, Table S10). *eCN+GCP-En1/2-tdT* CKO cells in both *Tbr1+* and *Olig2+* clusters showed altered expression of genes compared to their control counterparts (Fig. S7A, B), however GO analysis identified significant terms only for the *Tbr1+* cluster (Fig. S7C, Table S11). GO analysis determined that genes downregulated in the *Tbr1*+ cluster (*Epbhb1, Gja1, Flrt2* and *Pcdh17,* Figure S7D) belonged to synaptic gene and cell adhesion categories, suggesting that *En1/2* loss has a role in promoting differentiation and maturation of eCNm.

### Molecular subpopulations of eCNm are differentially vulnerable to the loss of *En1/2*

Given that the eCNm of E14.5 *eCN+GCP-En1/2* CKOs have altered transcriptomes related to neural maturation before some mutant cells begin to experience cell death, we next tested whether the six markers for molecular subpopulations of eCNm (Fig. 3,4) undergo changes in their domains in mutants. IF analysis and quantification of eCN in *eCNm-En1/2-tdT* CKO (Fig. 6, Fig. S8) and *eCN+GCP-En1/2* CKO (Fig. S9) animals revealed that at E14.5 only the TBR2+ subpopulation was greatly reduced in the two *En1/2* CKOs compared to their respective controls (100% loss in *eCNm-En1/2-tdT* CKO and 73.38% reduction in *eCN+GCP-En1/2* CKOs compared to controls)(Fig. 6A-K, Fig. S8A-I, Fig. S9A, B). This result is despite no reduction in overall eCN number at E14.5, and the other subpopulation markers appearing normal, except for a slight increase in the BARHL1+ eCN in *eCN+GCP-En1/2* CKOs.

**Figure 6:**
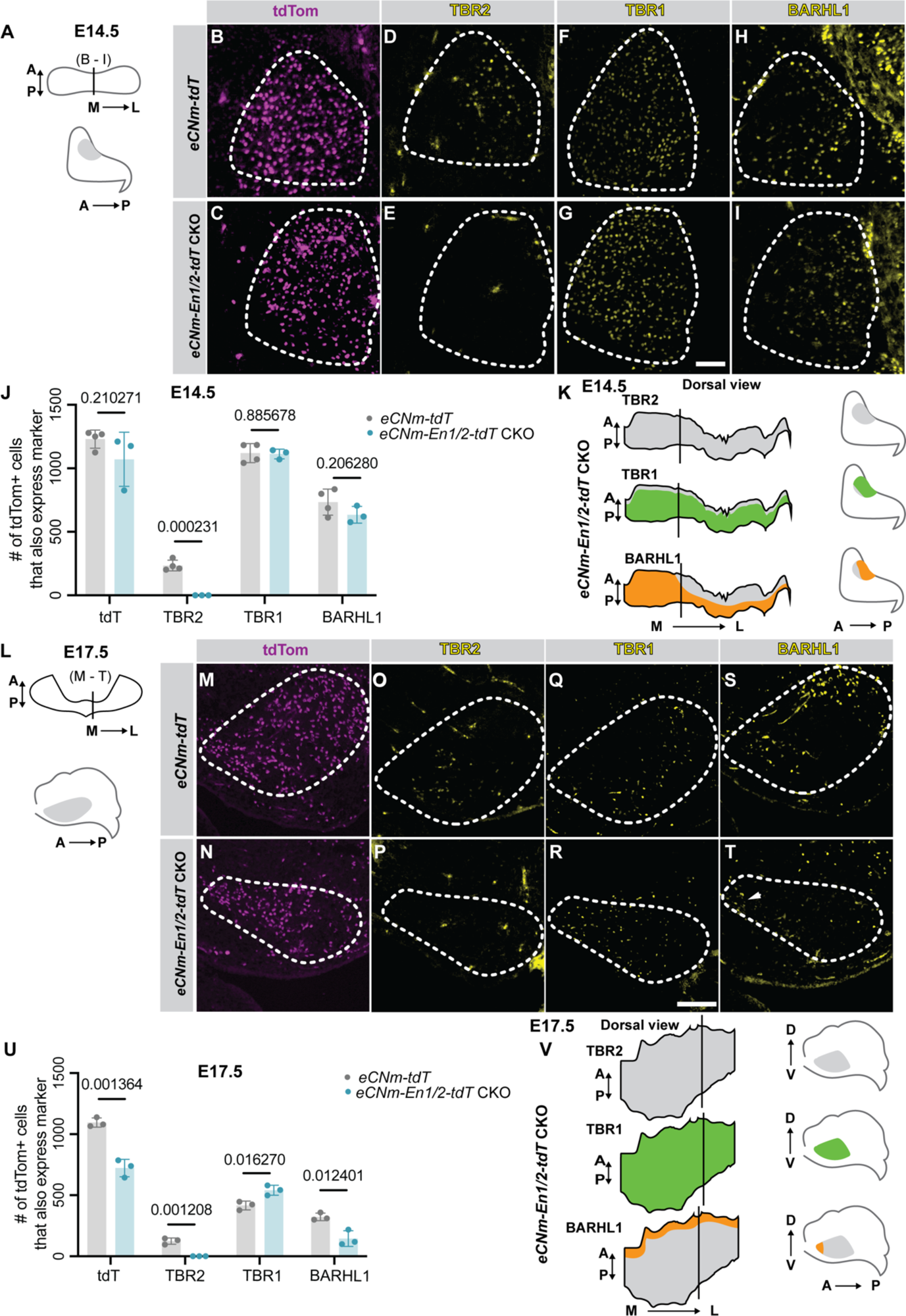
*En1/2* loss preferentially affects BARHL1, TBR2 and TBR1 eCNm expression domains. (A) Schematic of E14.5 cerebellum, with line indicating level of images in (B - I). (B - I) IF images of tdT (B,C), TBR2 (D,E), TBR1 (F,G) and BARHL1 (H,I) eCNm in E14.5 *eCNm-En1/2-tdT* CKO and *eCNm-tdT* control cerebella. (J) Quantification of tdT+ eCNm and TBR2, BARHL1 and TBR1 eCNm subpopulations in E14.5 *eCNm-tdT* and *eCNm-En1/2-tdT* CKO embryos (n = 4 *eCNm-tdT* and 3 *eCNm-En1/2-tdT* CKO animals: Student’s t-test). (K) Schematics of marker expression in eCNm of *eCNm-En1/2-tdT* CKO embryos projected onto 3D-reconstructions of the E14.5 *En1/2* CKO eCNm. (L) Schematic of E17.5 cerebellum, with line indicating level of images in (M - T). (M-T) IF images of tdT (M, N), TBR2 (O,P), TBR1 (Q,R) and BARHL1 (S,T) in E17.5 *eCNm-En1/2-tdT* CKO and *eCNm-tdT* cerebella. (U) Quantification of tdT+ eCNm and TBR2, BARHL1 and TBR1 eCNm subpopulations in E17.5 *eCNm-tdT* and *eCNm-En1/2-tdT* CKO embryos (n = 3 animals per genotype, Student’s t-test). (V) Schematics of marker expression in eCNm of *eCNm-En1/2-tdT* CKO embryos projected onto 3D-reconstructions of the E17.5 *En1/2* CKO eCNm. Dashed outlines: eCNm. Scale bars: 50 μm (B-I), 100 μm (M-T).

We then examined whether specific eCNm molecular subpopulations were lost or altered in their pattern at E17.5 given our previous finding that ∼18% of all eCN are lost to cell death in *eCN+GCP-En1/2* CKOs by this stage (Willett et al., 2019). As expected, the *eCNm-En1/2-tdT* CKO mutants had a significant reduction in tdT+ eCNm compared to controls (33.99%) (Fig. 6L-V). We then analyzed eCNm molecular subpopulations at E17.5 in the two *En1/2* CKOs to determine if specific subpopulations were susceptible to cell death and/or had altered gene expression compared to controls (Fig. 6O-V, Fig. S8J-R’, Fig. S9C, D). TBR2 was virtually absent in both 17.5 *En1/2* CKOs (100% reduction in *eCNm-En1/2-tdT* CKOs and 98.29% reduction in *eCN+GCP-En1/2-tdT* CKOs) (Fig. 6L-P, U; Fig. S9C, D). At E17.5, the loss of TBR2+ eCNm was likely due to a failure of eCNm to express TBR2 in the absence of *En1/2,* rather than cell death, as TBR1, which is normally co-expressed with TBR2+ in several anterior eCNm cells (Fig. S10), is not reduced in number and is in fact significantly increased in the two *En1/2* CKOs (Fig. 6Q, R, U; Fig. S9C, D). We also found that the BARHL1+ cells in the posterior-most eCNm were significantly reduced throughout the mediolateral axis of the E17.5 *En1/2* CKOs compared to their respective controls (total loss: 55.11% reduction in *eCNm-En1/2-tdT* CKO; 51.06% reduction in *eCN+GCP-En1/2-tdT* CKO compared to controls) (Fig. 6S-U, Fig. S9C, D). The only BARHL1+ eCNm remaining in two *En1/2* CKOs were the scattered cells in the anterior region. The domains marked by CALB2, PAX6 and LMX1A were unaffected both in pattern and number of eCNm cells in both *En1/2 CKOs* at both ages compared to controls (Fig. S8J-R, Fig. S9C,D). Thus, our spatial expression results show that embryonic eCNm subpopulations are differentially vulnerable to the loss of *En1/2,* and that the *En1/2* CKOs allow us to make a link between embryonic eCNm subpopulations and their adult counterparts (summarized in Fig. 7).

**Figure 7:**
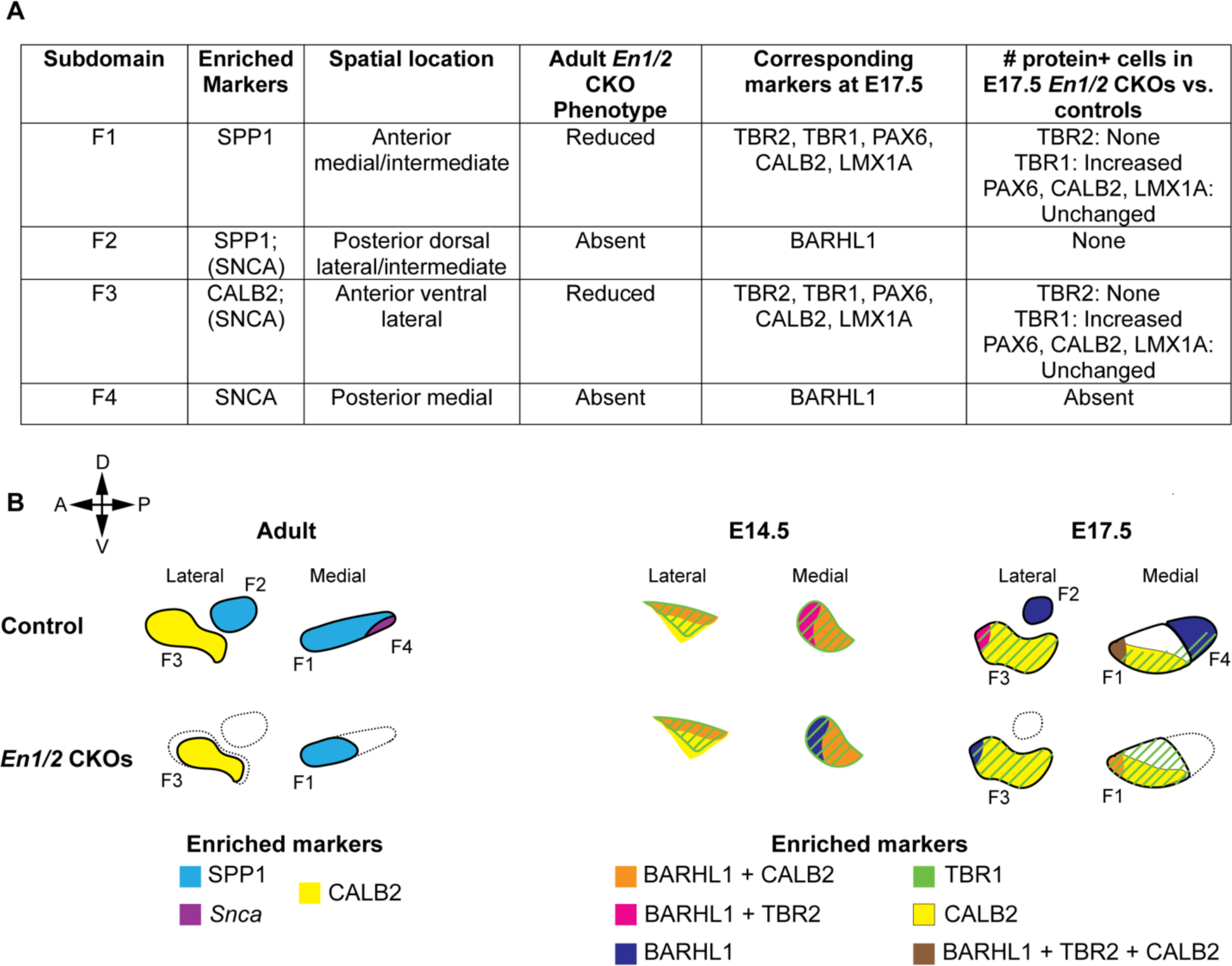
Proposed adult and embryonic eCNm subdomains, and alterations seen in *En1/2* conditional mutants. (A) Table listing proposed eCNm subdomains based on marker expression and spatial location in normal adult brain (based on Fujita et al, 2022) and at E17.5 in normal and *En1/2* CKOs (this study). Note, most proteins are expressed in a subset of cells in the four domains. Brackets around SNCA indicate where we could not detect expression. (B) Schematics of eCNm in the adult (left) and NTZ (E14.5) or eCNm (E17.5) embryos showing how the expression of some spatially defined marker proteins evolves during development and which subdomains are altered in *En1/2* CKOs. Solid black lines indicate F1-4 subdomains; dotted black lines indicate where control subdomain would be.

### TBR2 expression in GCPs and UBCs is dependent on *En1/2* function

Given that one of the earliest phenotypes of *En1/2* loss in the eCNm is a pronounced reduction of TBR2 expression, we asked whether the dependency of TBR2 on EN1/2 for its expression is recapitulated in the other two excitatory cerebellar cell types. Interestingly, we found that TBR2 is normally expressed in a subset of GCPs at E14.5 and E17.5 located toward the inner edge of the EGL (Fig. S11). TBR2 is also expressed in all UBCs from their birth to adulthood and these cells reside in the adult internal granule cell layer (IGL) and are enriched in posterior lobules 9 and 10 of the vermis (Englund et al., 2006).

We first determined whether TBR2 expression was lost in *En1/2* deficient UBCs at E17.5 in *eCNm-En1/2-tdT* CKO animals (Fig. S12), as the *eCN+GCP-Cre* transgene does not recombine in UBCs (Fig. S13). Strikingly, we found that TBR2 expression in UBCs at E17.5 was greatly reduced in *eCNm-En1/2-tdT* CKOs, and that in adults TBR2+ UBCs were dramatically reduced in number in the IGL compared to their respective controls (Fig. 8A-E, S12A-F). There were also fewer CALB2+ UBCs (marker of mature Type 1 TBR2+ UBCs) in adult *eCNm-En1/2-tdT* CKO mice compared to controls (Fig. 8F-J). Most of the tdT+ TBR2+/CALB2+ cells (UBCs) present in adult mutants were ectopically positioned outside the IGL in the white matter of lobule 10 or close to the CN, and any UBCs in the IGL appeared smaller than normal (Fig. 8K) and were concentrated in the ventral portion of lobule 10 and absent from lobule 9. These results indicate that while some UBCs can survive in the absence of *En1/2,* they are unable to mature normally or migrate to their proper position, a phenotype similar to that reported at birth for *Tbr2* null mutants (McDonough et al., 2021).

**Figure 8:**
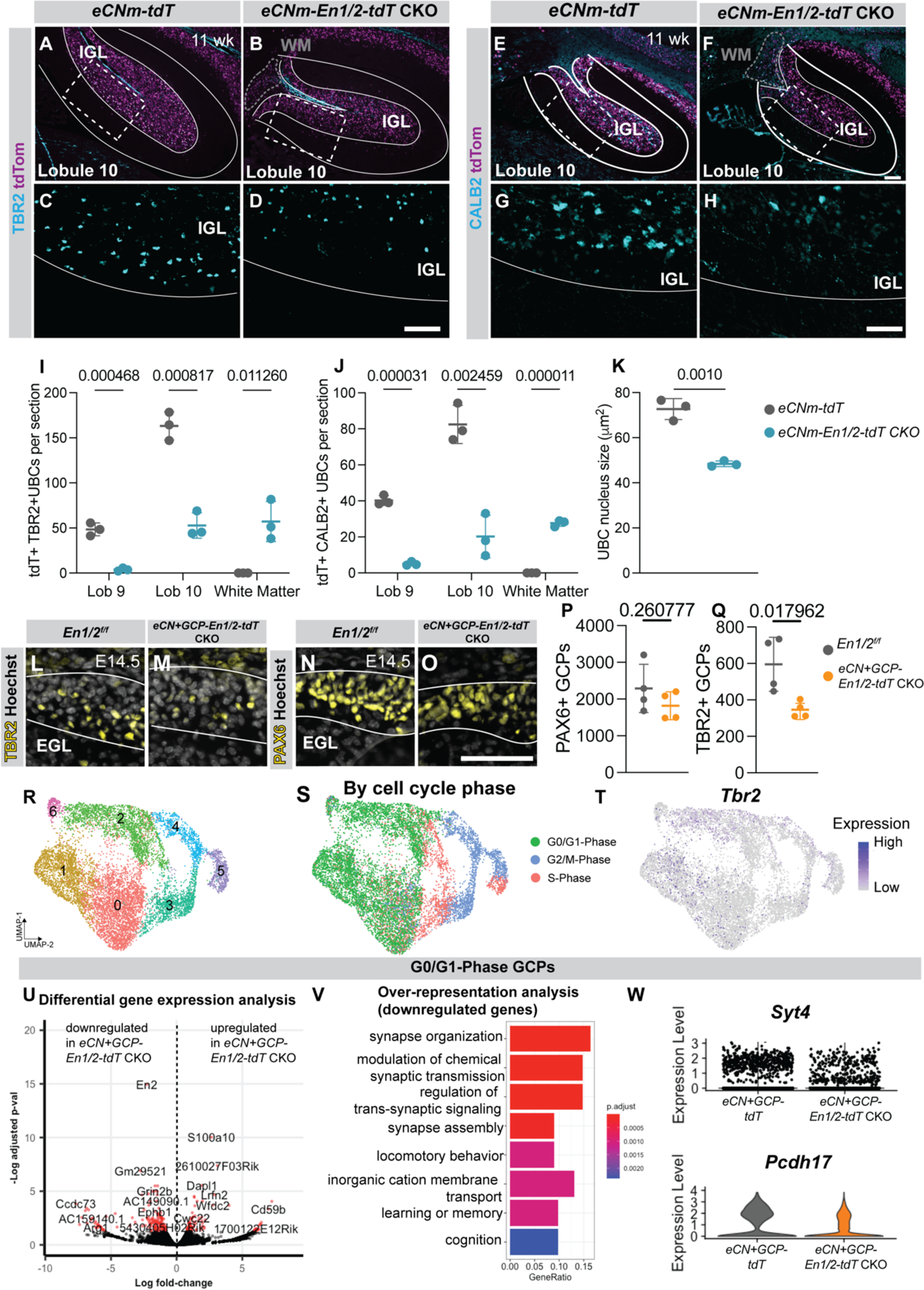
Deletion of *En1/2* results in reduction of TBR2 expression and differentiation defects in RL-derived GCPs and UBCs. IF images of TBR2 and tdT (A-D) or CALB2 and tdT (E-H) in lobule 10 of 11 week cerebella. Dashed rectangles (A,B,E,F) indicate regions from which images in (C,D,G,H) were taken. Quantification of tdT+ TBR2+ (I) or tdT+ CALB2+ (J) UBCs in *eCNm-En1/2-tdT* CKOs and *eCNm-tdT* controls in indicated lobules (Lob) or white matter (WM) and UBC nucleus size in lobule 10 (K) (n = 3 animals per genotype, Student’s t-test). IF images of TBR2+ (L,M) or PAX6+ (N,O) GCPs in EGL (white outlines) of E14.5 *En1/2^f/f^* and *eCN+GCP-En1/2-tdT* CKO cerebella. Quantification of PAX6+ GCPs (P) and TBR2+ GCPs (Q) (n = 4 per genotype, Student’s t-test) at E14.5 in two genotypes. (R) UMAP visualization of reclustering of *Pax6* enriched clusters shown in Fig 2A. (S) UMAP visualization with cells grouped by cell cycle phase. (T) *Tbr2* expression. (U) Volcano plot showing differentially expressed genes in G0/G1-Phase GCPs between two genotypes. Red dots indicate genes significantly up- or down-regulated (*P* ≤ 0.05), black dots indicate non-significantly enriched genes. (V) Over-representation analysis of downregulated genes in G0/G1-Phase GCPs of mutants compared to controls. (W) Violin plots showing expression of synaptic genes in G0/G1-Phase GCPs of both genotypes. Scale bars: 50 μm (A-D), (F-I), (L-O).

Our previous analyses on GCPs during the first postnatal week showed a mild diminution of differentiation in *En1/2* deficient GCPs compared to controls by analyzing mosaic mutants in which *En1/2* were deleted at P2 and analyzed at P8 (Willett et al., 2019). Interestingly, on sections of E14.5 and E17.5 *eCN+GCP-En1/2-tdT* CKO cerebella, few TBR2+ PAX6+ GCPs were detected despite PAX6 cell number being unaltered (Fig. 8L-Q and Fig. S14). We then combined the *Pax6+* clusters of E14.5 scRNA-seq tdT+ cells from *eCN+GCP-En1/2-tdT* CKO and control embryos and then reclustered the cells, which yielded 7 clusters which were partially distinguished by their cell cycle stage rather than by spatial markers (Fig. 8R-T). When we grouped GCPs by their cell cycle stage and performed differential gene expression analysis between *eCN+GCP-En1/2-tdT* CKOs and controls, over-representation analyses of downregulated genes in the G0/G1 phase *En1/2* CKO GCPs revealed that similar to *En1/2* deficient eCNm, *En1/2* deficient GCPs have a significant downregulation of genes and GO terms related to synapse formation and cell-cell adhesion (Fig. 8U-W, Table S12, S13). Taken together, these results show that EN1/2 positively regulate TBR2 expression in all cerebellar rhombic lip lineages. However, the severity of the consequences of *En1/2* loss and subsequent downstream TBR2 loss varies by cell type. Interestingly, by analyzing conditional mutants we found that UBCs are the most dependent on *Tbr2,* since *eCN+GCP-Tbr2* CKO mice (*Atoh1-Cre/+; Tbr2^lox/lox^*) do not have defects in eCN or granule cell survival (Fig. S15).

## DISCUSSION

In this study, we uncovered the underpinnings of embryonic eCNm subpopulation molecular diversity and defined overlapping and distinct roles for EN1/2 in all three cerebellar RL lineages. We first showed that the molecular identity of the posteriorly located subpopulation of the eCNm lost in *En1/2* CKOs (Willett et al, 2019 and Fig. 1, S1) is defined by *Snca* expression medially and SPP1 expression laterally relating to two of the adult eCNm domains recently described (F2, 4 in Fujita et al., 2020) (Figure 7). scRNA-seq and spatial protein expression analysis in a transgenic mouse *Cre* line that labels the eCNm at E14.5 revealed that the transcriptome of the eCNm is already distinct from the eCNi+eCNl one day after eCN neurogenesis is complete. Additionally, we define eCNm molecular subpopulations based on expression of marker proteins restricted to spatial domains that evolve between E14.5 and E17.5 (Fig. 7). scRNA-seq analysis of *En1/2* CKO eCNm indicates a role for EN1/2 in promoting neural differentiation, particularly synaptogenesis. Furthermore, eCNm subpopulations are differentially vulnerable to loss of *En1/2,* with a TBR2+ anterior eCNm subpopulation dependent on EN1/2 for their molecular identity, whereas a posterior BARHL1+ subpopulation relies on EN1/2 for viability (Fig. 7). Revealing global roles for EN1/2 in cerebellar excitatory neurons, we show that TBR2 expression and neural differentiation in GCPs and UBCs are also dependent on *En1/2,* with UBCs also requiring *En1/*2 for viability.

While molecular subpopulations of the adult eCN have been documented (Fujita et al,2020; Kebschull et al, 2020), how the populations emerge during development has not been understood. Complicating studies to draw direct connections between the embryonic and adult eCNm subpopulations, two of the three “steady state” markers of adult eCNm subpopulations are not similarly represented at embryonic stages (Fig. S3); SPP1 is not expressed in the embryo and *Snca* is broadly expressed. Be defining four molecular subdomains of the embryonic eCNm and determining which are lost in *En1/2* CKOs, we mapped two posterior E17.5 subpopulations of eCNm labeled by BARHL1 to the adult posterior domains expressing SPP1 (F2) and *Snca* (F4). In addition, a distinct anterior eCNm domain marked by CALB2 in the embryo and the adult (F3) is not lost in the *En1/2* CKOs. Interestingly, the anterior eCNm subdomains (F1 and F3) reduced in number in adult *En1/2* CKOs are not decreased in number at E17.5 (Fig. 7). The anterior eCNm thus appear to become sensitive to loss of *En1/2* for their viability at a later time point than the poster domains. Our results thus reveal that acquisition of subpopulation identity and vulnerability to *En1/2* loss occurs progressively soon after the eCNm reach the NTZ. However, the expression domains of genes that determine a particular subpopulation evolve between E14.5 and 17.5 and most embryonic genes are not expressed in the adult eCNm and thus the transcriptome of eCNm must continue to be refined after birth.

Our study highlights that transcription factors can play multiple roles through the development of a cell type. We show that *En1/2* are initially responsible in the E14.5 eCNm for spatial patterning (in a TBR2+ anterior domain) and enhancing synaptic gene expression, followed by a role in survival (in BARHL1+ posterior domains). Similar roles have been documented for the *engrailed* genes in several species. For example, in the fly, *en* is responsible for directing CNS midline glia to a posterior, non-axon ensheathing fate by repressing anterior, axon-ensheathing glial genes (Watson et al., 2011). Additionally, *en* in the fly and *En1/2* in the mouse are critical for the proper development of serotonergic neurons (Fox and Deneris, 2012; Lundell et al., 1996). In both species, the *engrailed* genes control expression of genes critical to serotonin synthesis, and in the mouse, loss of *En1/2* results in apoptosis of 60% of dorsal raphe nucleus neurons starting at postnatal day 10 (Fox and Deneris, 2012). However, whether the neurons that die belong to a molecularly defined subpopulation(s) remains to be determined. Additionally, *en* in the fly auditory system controls synapse formation and specificity of a subset of neurons (Pézier et al., 2014). In the mouse, *En1* is required in a subset of spinal cord interneurons to make the appropriate number of recurrent inhibitory connections to motor neurons (Sapir et al., 2004). How EN1/2 mediate transcriptional control over these processes, however, remains to be elucidated.

EN1/2 are thought to primarily act as repressors (Jaynes and O’Farrell, 1991; Smith and Jaynes, 1996; Tolkunova et al., 1998) and thus absence of *En1/2* would be expected to lead to upregulation of genes. Therefore, the downregulation of synaptic genes we observed in eCNm could be due to an indirect process via the repression of a repressor. However, en in the fly has been shown to act as an activator in the presence of its co-factor, extradenticle (PBX proteins in vertebrates) (Serrano and Maschat, 1998). Moreover, chromatin immunoprecipitation data of genes regulated by both en and gooseberry-neuro (PAX3/7 in vertebrates) in the ventral nerve cord of the fly, which act in concert to drive posterior commissure crossing, showed that most combined targets are activated, rather than repressed (Bonneaud et al., 2017). Most target genes belong to nervous system development GO categories, including axon guidance (Bonneaud et al., 2017). Thus, it is possible that EN1/2 in the mouse eCNm activate synaptic gene expression directly in the presence of unidentified partner transcription factors or co-factors.

We discovered that *En1/2* are required for the activation of TBR2 expression in all three RL-derived excitatory neurons, indicating one general mechanism by which *En1/2* likely controls neural differentiation. In the cerebral cortex TBR2 and TBR1 play sequential roles in neural differentiation and acquisition of laminar fate (Bedogni et al., 2010; Mihalas et al., 2016). Similar roles in controlling neural differentiation have been suggested for the proteins in the NTZ based of their expression patterns at E13.5 (Fink et al., 2006). However, our study shows that TBR1 marks the majority (∼80%) of eCNm at E14.5, while TBR2 is only expressed in ∼20% of the eCNm, suggesting TBR1 is the main player in eCNm development. Consistent with this, we found that conditional loss of *Tbr2* does not alter eCN or GC number, and adult cerebellar morphology and size appear normal (Fig. S15). However, these results do not rule out a critical role for the TBR family in eCNm development since TBR1 is present in most TBR2+ eCNm. In contrast, TBR2 plays a pivotal role in the differentiation, survival and migration of the UBCs where TBR1 is not expressed (Fig. S12) and TBR2 expression is maintained from the time the UBCs are born into adulthood (Fig. 8, S12, 13, Englund et al., 2006; McDonough et al., 2021). Strikingly *En1/2* deficient adult UBCs haver a similar phenotype to that reported for *Tbr2* deficient UBCs at birth (McDonough et al, 2021) a cell type where TBR1 is not upregulated in *En1/2* deficient UBCs as a possible means of compensation (Fig. S12). Curiously, TBR2 does not play a major role in development of GCPs, despite lack of upregulation of TBR1 expression in *En1/2* CKO GCPs (Fig. S13). Thus, EN1/2 and TBR2 have distinct roles in the three RL-derived excitatory neuron types, and EN1/2 are upstream of TBR2 but seemingly not *Tbr1,* or *Pax6* another homeobox gene with differential roles in the RL-derived lineages (Engelkamp et al., 1999b; Swanson and Goldowitz, 2011; Swanson et al., 2005).

In conclusion, our study sheds light on how molecularly defined neuron subpopulations emerge, and how transcription factors that initially pattern a large region of tissue early in development can be critical later in defining subpopulations of cells in a lineage and then for their differentiation and/or homeostasis.

## MATERIALS AND METHODS

### Animals

All animal experiments were performed in accordance with the protocols approved and guidelines provided by the Memorial Sloan Kettering Cancer Center’s Institutional Animal Care and Use Committee (IACUC). Animals were given access to food and water ad libitum and were housed on a 12 hr light/dark cycle. The following mouse lines were used in this study: *En1^lox^* (Sgaier et al., 2007), *En2^lox^*(Cheng et al., 2010), *Atoh1-Cre* (Matei et al., 2005), *R26^LSL-ntdTom^*(Quina et al., 2017)(Ai75D, Jackson Labs, stock no: 025106), *Selenow-Cre* (Gerfen et al., 2013) and *Tbr2^lox^* (Intlekofer et al., 2008). Primers used for genotyping are listed in Table S14. Animals were maintained on an outbred Swiss Webster background. Both sexes were used for all analyses and no randomization was used. Noon of the day a vaginal plug was detected was designated as developmental stage E0.5.

### Tissue processing

The brains of all embryonic stages were dissected in ice cold phosphate buffered saline without calcium and magnesium (PBS) and immersion fixed in 4% paraformaldehyde (PFA) for 48 hr (for E14.5) and 72 hr for E17.5 at 4°C on a shaker. Adult animals were anesthetized with isoflurane and then transcardially perfused with ice cold PBS followed by 4% PFA, and brains were post-fixed in 4% PFA overnight. Specimens for cryosectioning were placed in 30% sucrose in PBS until they sank, embedded in OCT (Tissue-Tek), frozen in dry ice cooled isopentane, and sectioned on a cryostat in the sagittal plane at 12 μm for embryos and 14 μm for adults (Leica, CM3050S) and collected on glass slides (Fisherbrand). Tissue was stored at -20°C.

### Immunofluorescence

Primary antibodies and their related concentrations are listed in Table S14. Sections of cryosectioned tissue were air dried for 10 minutes. They were then washed in PBS for 10 minutes. For immunofluorescence with all antibodies, sections were subjected to antigen retrieval using sodium citrate buffer (10 mM sodium citrate with 0.05% Tween-20, pH 6.0) at 95℃ for 15 minutes (embryonic tissue) or 20 minutes (adult tissue). After antigen retrieval, sections were brought to room temperature. Sodium citrate buffer was discarded and then sections underwent two washes in 0.1% PBST (1X PBS with 0.1% Triton-X) for 5 minutes each. Slides were then blocked with blocking buffer (5% Bovine Serum Albumin (Sigma) in 1X PBS with 0.1% Triton X-100) at room temperature for 1 hour. Then, primary antibodies diluted in blocking buffer were placed on slides overnight at 4℃. Slides were washed three times in 0.1% PBST for 5 minutes each and then incubated in secondary antibodies (Alexa Fluor conjugated secondary antibodies) diluted at 1:500 in blocking buffer for 1 hour at room temperature. Counterstaining was performed using Hoechst 33258 (1:1000, Invitrogen A-21422). Slides were then washed three times in PBS for 5 minutes each prior to cover slipping using Fluorogel mounting medium (Electron Microscopy Sciences).

### RNA in situ hybridization

Probes were in vitro transcribed from PCR-amplified templates prepared from cDNA synthesized from postnatal cerebellum lysate. Primers used for PCR amplification are listed in Table S14. Primers were flanked in the 5’ with SP6 (antisense) and T7 (sense) promoters. Specimen treatment and hybridization were performed as described previously (Blaess et al., 2011).

### Hematoxylin and Eosin Staining

Hematoxylin and Eosin (H & E) staining (Thermofisher) was performed according to instructions from the manufacturer for histological analysis and cerebellar area measurements of adult sections.

### Nickel enhanced DAB Staining

eCN counting for adult *eCN+GCP-Tbr2* CKOs was performed using nickel enhanced DAB (Ni-DAB) immunohistochemistry for NeuN as previously described (Willet et al., 2019).

### Microscopy

All images were collected with a DM6000 Leica fluorescent microscope or NanoZoomer Digital Pathology microscope (Hamamatsu Photonics) and processed using ImageJ (NIH) or Photoshop (Adobe) software. Image quantification, including area measurements and cell counting was performed using ImageJ (NIH). The polygon selection tool (ImageJ) was used to select areas of interest for all cerebellar area measurements, and the Cell Counter plugin (ImageJ) was used to quantify all cells (aside from adult eCN).

### Embryonic eCN and GCP quantifications

For all embryonic eCN and GCP quantifications, every tenth section from a hemisphere section where the CN first appeared to the midline was quantified. Cells were only quantified if they expressed both the marker and tdT.

### UBC quantifications

For adult UBC quantifications, an average of tdT+ and TBR2+ or CALB2+ cells was calculated from quantifying lobules 9 and 10 in three sections per mouse. Nuclei were measured using the “oval” tool on ImageJ. 33 or 34 cells were measured from each section, either from lobule 10 or the adjacent white matter near the CN (for *eCNm-En1/2-tdT* CKOs).

### Adult eCN quantifications

Adult eCN quantifications were performed using NeuN labeled slides (every other section for *eCN+GCP-Tbr2* CKOs) using a semi-automated method described in Willett et al., 2019. Briefly, eCN from scanned images (Nanozoomer Digital Pathology) were cropped, pre-processed in ImageJ and only particles sized 100–600 um^2^ were quantified.

### Adult cerebellar area measurements

For all adult sector and IGL areas, H & E stained slides were used. Three sagittal sections at midline per animal were used, and values across these sections were averaged.

### Three-dimensional (3D) reconstructions

Every other section (12 microns thick) from a consecutive sagittal series was used for 3D reconstruction from where the eCN appeared laterally to where they disappeared medially. eCNm were cropped in Adobe Photoshop, and then imported into ImageJ. The images were registered using the StackReg plug-in on ImageJ. Minor adjustments were made manually only where needed. Registered images were imported into Adobe Illustrator and each image was assigned its own layer. Each region consisting of the eCNm, marked by tdT, was outlined in Adobe Illustrator. Aligned outlines were imported into Rhinoceros 7 and were spaced evenly by distance between sections using the SetPt command. The final 3D reconstruction was made using the Loft command (parameters: 20 control points, loose). Dorsal views of rendered images were outlined in Adobe Illustrator (using the Image Trace function) and colored for schematics.

### Statistical analyses

Prism (GraphPad) was used for all statistical analyses except for genomics analyses. Statistical comparisons of two populations (genotypes) were done with Student’s two-tailed t-test and Two-way ANOVA. *Post hoc* analyses of Two-way ANOVA was performed using Šídák multipe comparison tests. The statistical significance cutoff was set at *P* ≤ 0.05. Population statistics were represented either as mean ± standard deviation (s.d) of the mean or as mean ± standard error of mean (s.e.m). No statistical methods were used to predetermine the sample size, but our sample sizes are similar to those generally employed in the field. n ≥ 3 mice were used for each experiment and the numbers for animals used for each experiment are stated in the figure legends.

### Sample preparation for single cell RNA-sequencing

Animals (numbers of embryos pooled for each genotype are listed in relevant figures) were dissected in ice cold Hank’s balanced salt solution (HBSS) (Gibco) and were pooled together for downstream steps. Two replicate experiments were performed. On ice, cerebella were detached from the rest of the brain using forceps and dissociated in Accutase (Innovative Cell Technologies) at 37°C for 7-10 minutes. Following dissociation, accutase was washed out using neural stem cell (NSC) media (Neurobasal, supplemented with N2, B27 and non-essential amino acids, Gibco). Samples were triturated by pipetting up and down using a P1000, filtered through a 35um cell strainer (Falcon) and then centrifuged at 500g at 4°C for 5 minutes. Cells were washed again in NSC media and were finally filtered into polypropylene round bottom tubes (Falcon) prior to sorting for tdT+ cells. tdT+ cells were sorted into polypropylene round bottom tubes, spun down at 500g at 4°C for 5 minutes and resuspended in NSC media prior to single cell library preparation.

### Library preparation and sequencing

Single cell libraries were prepared using 10X Genomics’ Chromium Next GEM Single Cell 3’ Reagent Kits, v3.1 according to manufacturer’s instructions. Prior to loading the 10X Genomics chip, the limited sample preparation protocol was used. Cell viability was quantified using 0.2% (w/v) Trypan Blue on a hemocytometer, and 7,000 viable cells were targeted per pooled sample.

After PicoGreen quantification and quality control by Agilent TapeStation, libraries were pooled equimolar and run on a NovaSeq 6000 in a PE28/91 run, using the NovaSeq 6000 S1 Reagent Kit (100 Cycles) (Illumina).

### Single cell RNA sequencing analyses

The Cell Ranger Single Cell software suite from 10x Genomics was used to align reads and generate feature-barcode matrices. The reference genome used was the Genome Reference Consortium Mouse Build 38 (mm10). Raw reads were processed using the Cell Ranger count program using default parameters.

Seurat v4.1.1 was used to generate a UMI (unique molecular identifier) count matrix from the Cell Ranger output (Butler et al., 2018; Macosko et al., 2015). Genes expressed in less than 10 cells were removed from further analyses. Cells that had fewer than 500 UMIs, 250 genes or had over 30% of their UMIs mapped to mitochondrial genes were considered low quality/outliers and discarded from the datasets. Normalization was performed on individual samples using the scTransform function, with regression of *Xist.* Samples were then integrated using the PrepSCTIntegration, FindIntegrationAnchors and the IntegrateData functions. For all analyses, the first 25 dimensions were used for the FindNeighbors function, and clusters were identified using the FindClusters function with a resolution of 0.5. Data were projected into the 2D space using the FindUMAP function with 25 dimensions. Cluster markers and further differential gene expression analyses were all performed on the RNA assay, following normalization using the NormalizeData functions. Cluster markers were identified using the FindAllMarkers function (for *eCN+GCP-tdT* or *eCN+GCP-En1/2-tdT* CKO only analyses) or FindConserved markers (for joint control and *eCN+GCP*-*En1/2-tdT* CKO analyses) and comparing markers generated to existing literature on the embryonic cerebellum and analysis of the Allen Developing Brain Expression Atlas. To refine clustering further, the SubsetData function was used to create a new Seurat object, and the above clustering was reiterated. For the *eCN+GCP-En1/2-tdT* CKO only clustering, selection of clusters for downstream reclustering as well as final mapping of the eCN clusters was performed using the FindTransferAnchors and TransferData functions with the corresponding *eCN+GCP-tdT* dataset as the reference dataset.

Differential gene expression analysis between controls and *En1/2* CKOs was performed using Libra (Squair et al., 2021) using the edgeR-LRT pseudobulk method. Genes that showed an adjusted p-value σ≤ 0.05 were considered significantly up or downregulated. Results were visualized using the EnhancedVolcano package on Bioconductor. Gene ontology analyses were performed separately on up and downregulated genes using the clusterProfiler (Wu et al., 2021; Yu et al., 2012) package on Bioconductor.

## ACKNOWLEDGEMENTS

We thank past and present members of the Joyner lab for helpful comments throughout the work. We also thank the Memorial Sloan Kettering Cancer Center Flow Cytometry Core, Integrated Genomics Operations Core Facility and the Center for Comparative Medicine and Pathology for technical support. We acknowledge the use of the Integrated Genomics Operation Core, funded by an NCI Cancer Center Support Grant (CCSG, P30 CA08748), Cycle for Survival, and the Marie-Josée and Henry R. Kravis Center for Molecular Oncology. We are grateful to Natalia De Marco Garcia for the *Selenow-Cre* mice and to Joseph Sun for the *Tbr2^lox^* mice. We thank Thomas Vierbuchen for valuable feedback and Daniel Medina Cano for assistance with using the 10X Chromium Controller. We thank Armita R. Manafzadeh for assistance with 3D reconstruction of the eCNm.

## COMPETING INTERESTS

The authors declare no competing or financial interests.

## FUNDING

This work was supported by grants from the NIH to ALJ (R37MH085726 and R01NS092096) and ASL (NICHD T32HD060600) and a National Cancer Institute Cancer Center Support Grant [P30 CA008748-48]. The work was also supported by Cycle for Survival funds. AK was supported by a Dorris J. Hutchison Pre-doctoral Fellowship. ASL was also supported by a Weill Cornell Medicine Clinical & Translational Science Center Predoctoral Training Award (TL1TR002386) from National Center for Advancing Translational Sciences. NSB was supported by postdoctoral fellowships from NYSTEM (C32599GG) and NIH/NINDS (K99/R00 NS112605-01). ON was supported by the McNulty Scholars Program at Hunter College.

## DATA AVAILABILITY

scRNA-seq data are available as GEO accession GSE246894. Original 3D reconstruction files can be accessed at 10.6084/m9.figshare.24650286.

---

**Figure S1:**
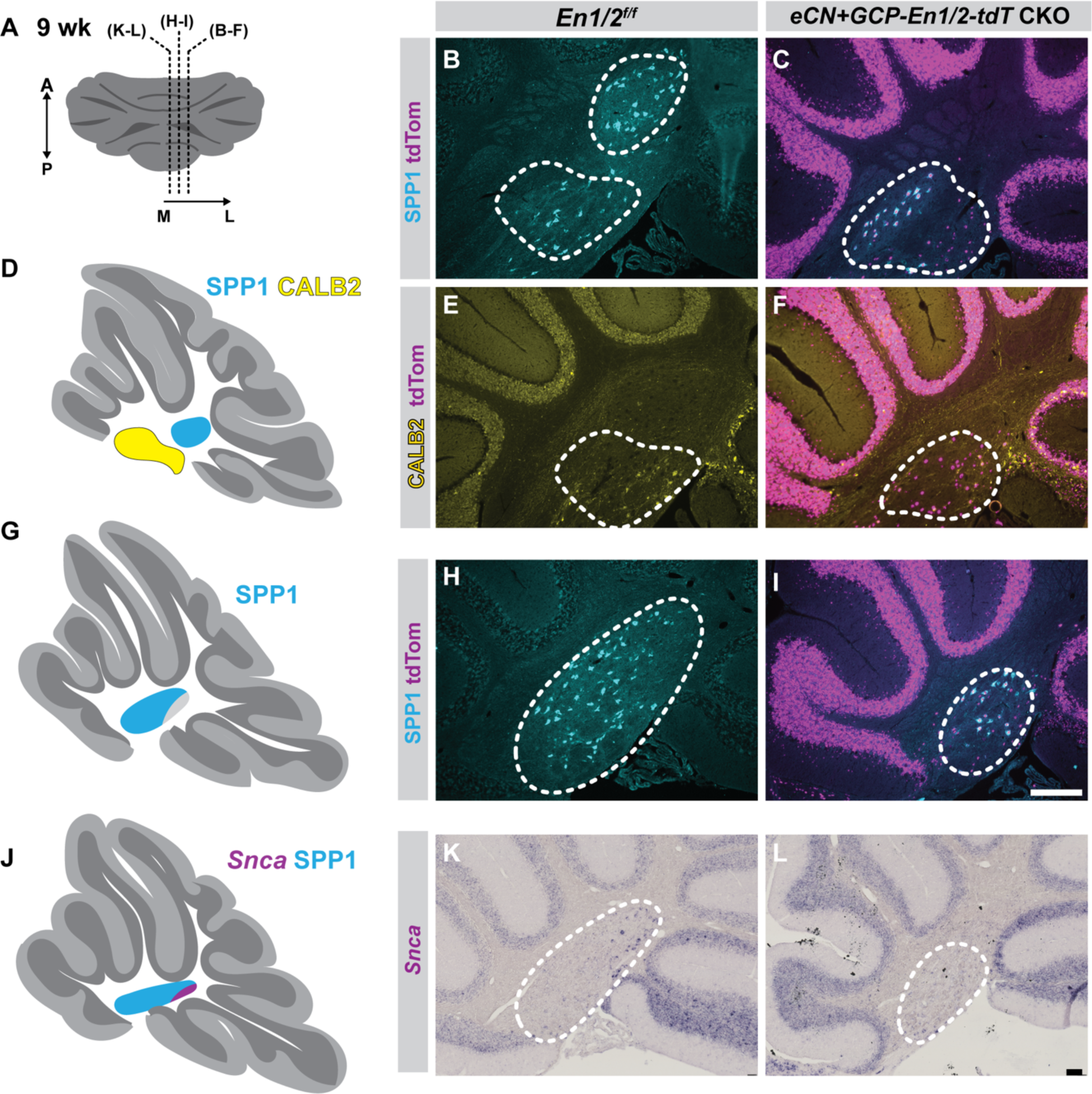
Two posterior subpopulations of adult eCNm are absent in *eCN+GCP-En1/2-tdT* CKO animals and anterior domains are diminished. (A) Schematic of cerebellum indicating levels at which images in (B-L) were taken. (B-F) Immunofluorescence (IF) analysis of SPP1 and CALB2 on sagittal sections of 9 week *eCN+GCP-En1/2-tdT* CKO and controls at a lateral level. (E-I) IF analysis of SPP1 at an intermediate level. (K-L) ISH analysis of *Snca* at a medial level. White dashed outlines indicate eCNm. Scale bars: 100 μm.

**Figure S2:**
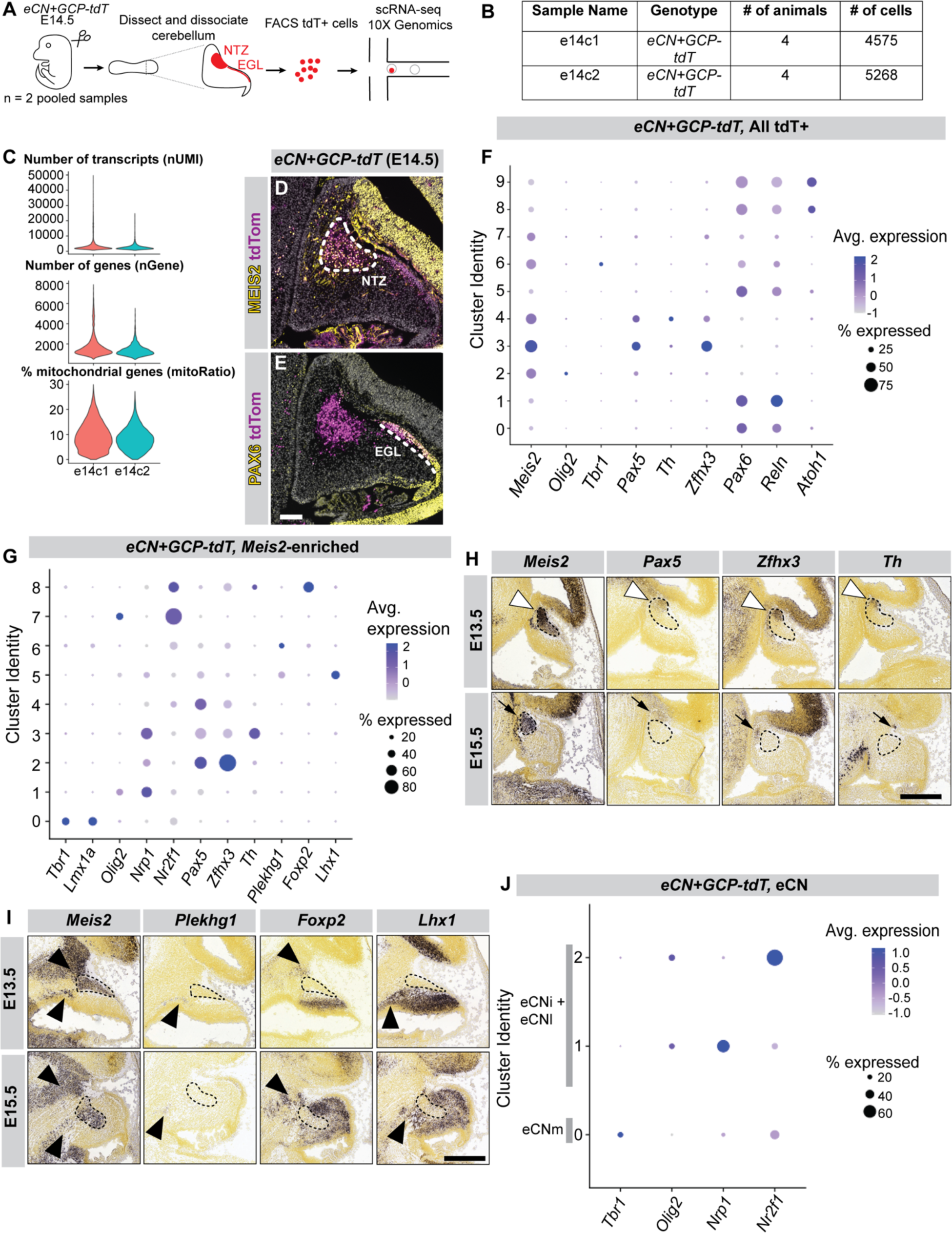
Quality control and marker analysis of tdT+ cells from *eCN+GCP-tdT* scRNA-seq dataset. (A) Schematic of scRNA-seq experiment. (B) Table detailing animal number and cell number per replicate. (C) Violin plots depicting number of transcripts (nUMI), number of genes (nGene) and percentage of mitochondrial genes (mitoRatio) by replicate. (D) IF image of sagittal section from E14.5 *eCN+GCP-tdT* cerebellum showing colocalization of MEIS2 and tdT in the NTZ (dashed outline). (E) IF image of sagittal section from E14.5 *eCN+GCP-tdT* cerebellum showing colocalization of PAX6 and tdT in GCPs in the EGL (dashed line). At this level PAX6 is only present in some eCNm. (F) Marker expression for all tdT+ clusters. (G) Marker expression for all *Meis2*-enriched clusters after reclustering. (H) RNA in situ hybridization (ISH) analysis of sagittal sections of E13.5 (top row) and E15.5 (bottom row) cerebella showing expression of marker genes in the anterior NTZ (empty arrowheads). Note *Pax5, Zfhx3, Th* (Meis2 enriched clusters 2, 3, 4) are expressed in the anterior NTZ at E13.5 that does not contain eCN (dashed outlines) at E15.5 but are present in isthmic regions (black arrows). (I) ISH images of sagittal sections of E13.5 (top row) and E15.5 (bottom row) cerebella showing extracerebellar cells expressing *Meis2* and marker genes (black arrowheads). Note *Lhx1, Plekhg1* and *Foxp2* (Meis2 enriched clusters 5,6, 8) are expressed outside the cerebellum. ISH images from Allen Developing Mouse Brain Atlas. Scale bars: 100 μm (D,E), 500 μm (H,I).

**Figure S3:**
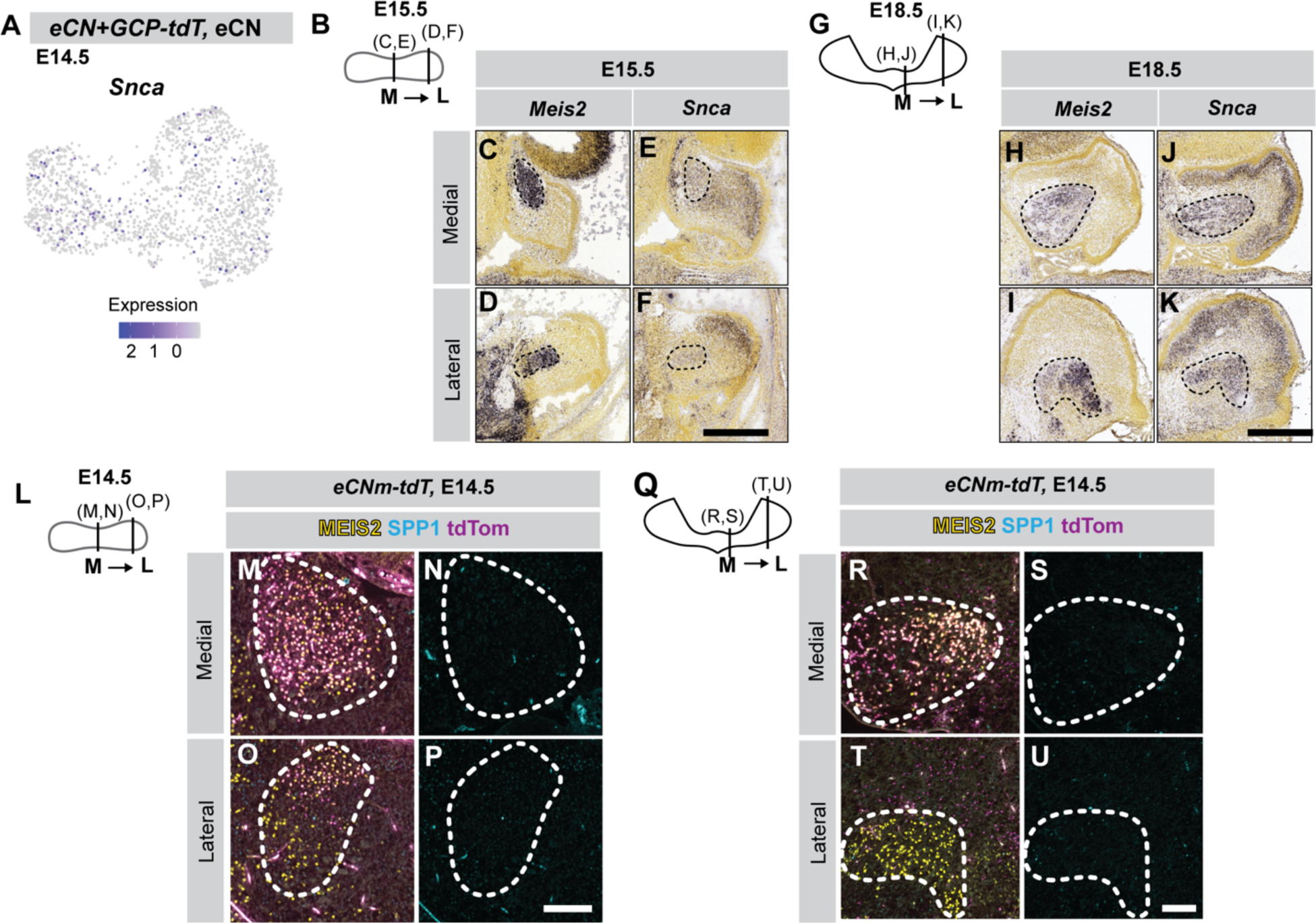
Adult eCNm subpopulation markers are not well represented in the E14.5 *eCN+GCP-tdT* eCNm scRNA-seq dataset and by section ISH analysis. (A) Expression of *Snca* in the eCN. (B) Schematic of E15.5 cerebellum indicating medial (C,E) and lateral (D,F) levels that sections were taken from. *Meis2* (C,D) and *Snca* (E,F) ISH images of E15.5 cerebellum. Note low expression throughout the eCN of *Snca* (dashed outlines). (G) Schematic of E18.5 cerebellum indicating levels at which (H,J) and (I,K) were taken from. *Meis2* (H,I) and *Snca* (J,K) ISH images of the E18.5 cerebellum. Note broad expression of *Snca* throughout the eCN (dashed outlines). ISH from the Allen Developing Mouse Brain Atlas. (L) Schematic of E14.5 cerebellum indicating levels at which images in (M,N) and (O,P) were taken. Merge of tdT and MEIS2 and SPP1 (M,O) and SPP1 single channel (N,P) with dashed line outlining the in E14.5 *eCNm-tdT* labeled cells. Note lack of SPP1 expression in the eCN. (Q) Schematic of E17.5 cerebellum indicating levels at which images in (R,S) and (T,U) were taken. Merge of tdT, MEIS2 and SPP1 (R,T) and SPP1 single channel (S,U) IF images of eCN (dashed outlines) in E17.5 *eCNm-tdT* mice showing no SPP1 expression in the eCN. SPP1 UMAP could not be generated as no SPP1 was detected in any cells from *eCN+GCP-tdT* scRNA-seq dataset. Scale bars: 500 μm (C-F), (H-K); 100 μm (M-P), (R-U).

**Figure S4:**
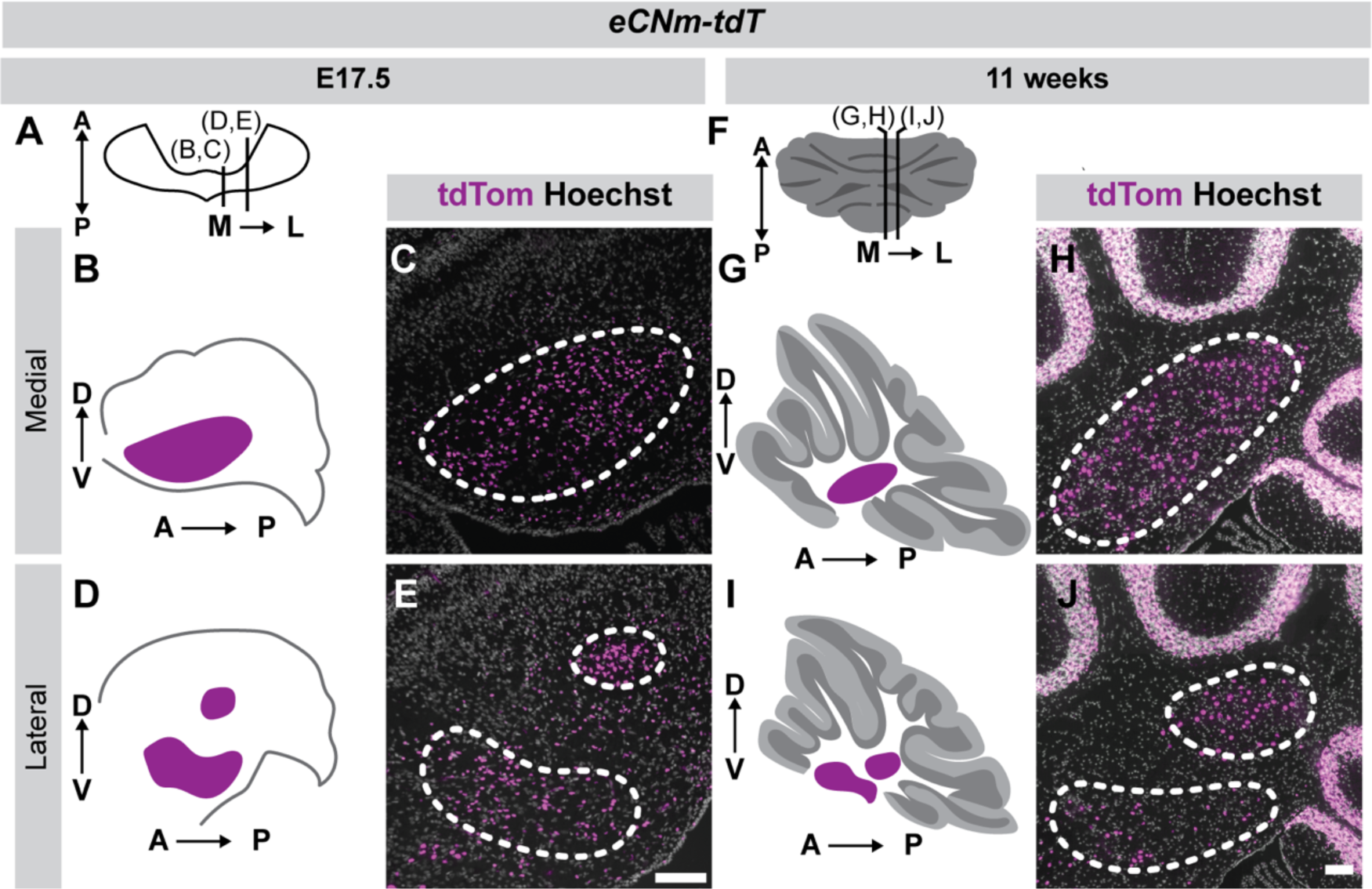
Morphology of eCNm is similar between E17.5 and adulthood. (A) Schematic of E17.5 cerebellum, with lines indicating mediolateral levels at which images in (B,C) and (D,E) were taken. (B) Schematic of sagittal section of the E17.5 cerebellum at a medial level of the eCNm (indicated in purple). (C) IF image of tdT+ eCNm (dashed line) in an E17.5 *eCNm-tdT* cerebellum at a medial eCNm level. (D) Schematic of sagittal section of the E17.5 cerebellum at a lateral level of the eCNm (indicated in purple). (E) IF image of tdT+ eCNm (dashed line) in an E17.5 *eCNm-tdT* cerebellum at a lateral eCNm level. (F) Schematic of an 11 week (adult) cerebellum, with lines indicating mediolateral levels at which images in (G,H) and (I,J) were taken. (G) Schematic of sagittal section of the adult cerebellum at a medial eCNm level (indicated in purple). (H) IF image of eCNm (dashed line) in the adult *eCNm-tdT* cerebellum at a medial eCNm level. (I) Schematic of sagittal section of the adult cerebellum at a lateral eCNm level (indicated in purple). (J) IF image of eCNm (dashed line) in an adult *eCNm-tdT* cerebellum at a lateral eCNm level. Note similarities between shapes of eCN in (C,H) and (E,J) respectively. Scale bars: 100 μm (C,H,E,J).

**Figure S5:**
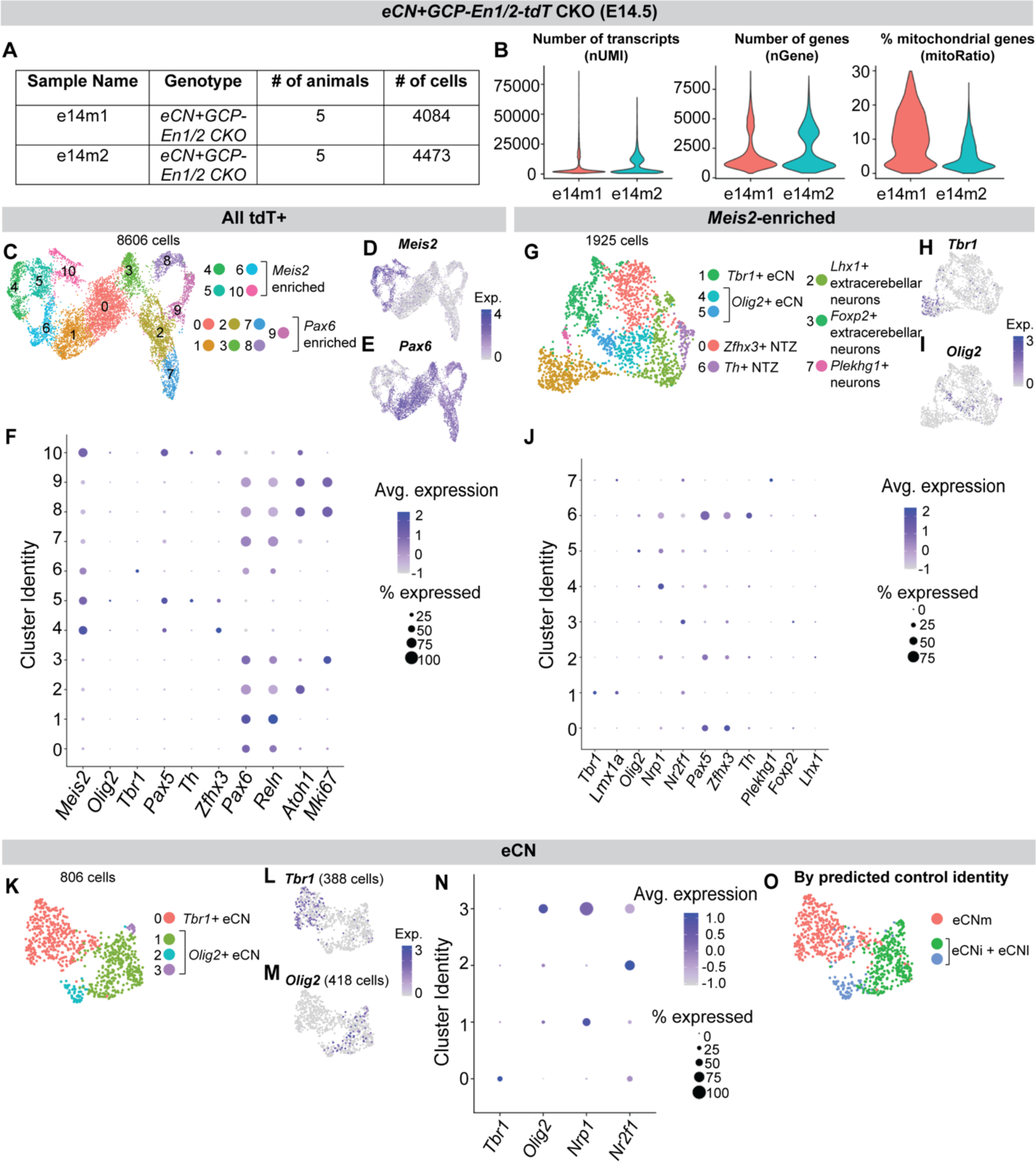
Quality control and marker analysis of tdT+ cells from *eCN+GCP-En1/2-tdT* CKO scRNA-seq dataset. (A) Table detailing mutant number and cell number per replicate. (B) Violin plots depicting nUMI, nGene and percentage of mitoRatio by replicate. (C) UMAP visualization of tdT+ cell clusters from *eCN+GCP-En1/2-tdT* CKO E14.5 cerebella, with *Meis2* (D) and *Pax6* (E) expression shown. (F) Select marker gene expression for all mutant tdT+ clusters. (G) UMAP visualization of *Meis2-*enriched mutant clusters after re-clustering, with *Tbr1* (H) and *Olig2* (I) expression shown. (J) Marker expression for all mutant *Meis2*-enriched clusters after reclustering. (K) UMAP visualization of reclustering of *Tbr1+* and *Olig2+* mutant clusters with *Tbr1* (L) and *Olig2* (M) expression shown. (N) Marker gene expression for all *Tbr1* and *Olig2* mutant clusters after reclustering. (O) Mutant eCN cluster cells colored by predicted control identity.

**Figure S6:**
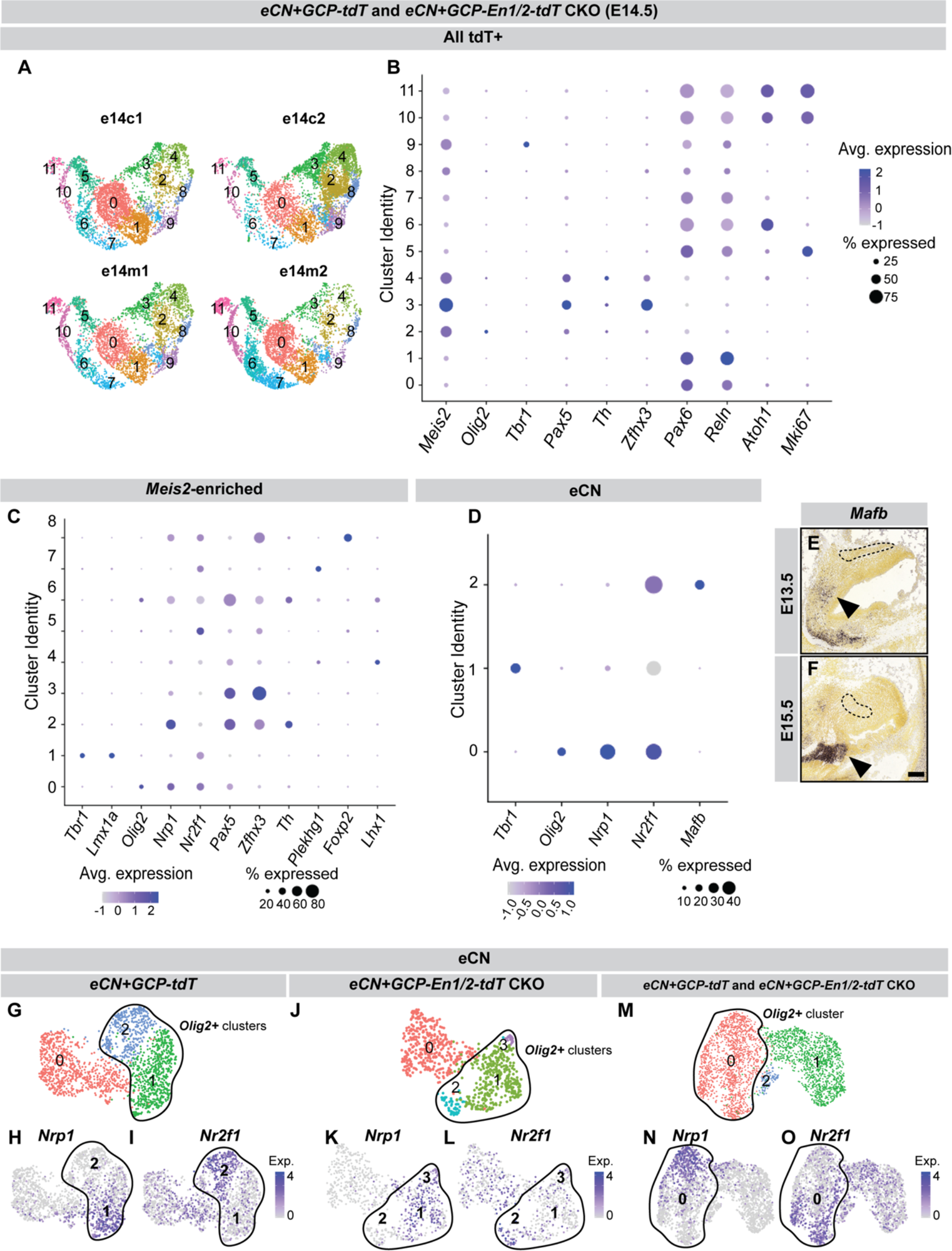
Marker analysis of scRNA-seq for *eCN+GCP-tdT* and *eCN+GCP-En1/2-tdT* CKO tdT+ cells combined. (A) Distribution of tdT+ cells across all clusters of mutant and control cells combined (m+c) by replicate. (B) Marker expression for all m+c tdT+ clusters. (C) Marker expression for all *Meis2*-enriched m+c clusters after reclustering. (D) Marker expression for all *Tbr1* and *Olig2*-enriched m+c clusters after reclustering. (E, F) ISH images of sagittal sections of E13.5 and E15.5 cerebellar showing expression of *Mafb* (from Allen Developing Mouse Brain Atlas). (G-I) UMAP of eCN clustering from *eCN+GCP-tdT* control samples, with expression of *Nrp1* and *Nr2f1.* (J-L) UMAP of eCN clustering from *eCN+GCP-En1/2-tdT* CKO mutant samples, with expression of *Nrp1* and *Nr2f1.* (M-O) UMAP of eCN clustering from m+c samples, with expression of *Nrp1* and *Nr2f1*.

**Figure S7:**
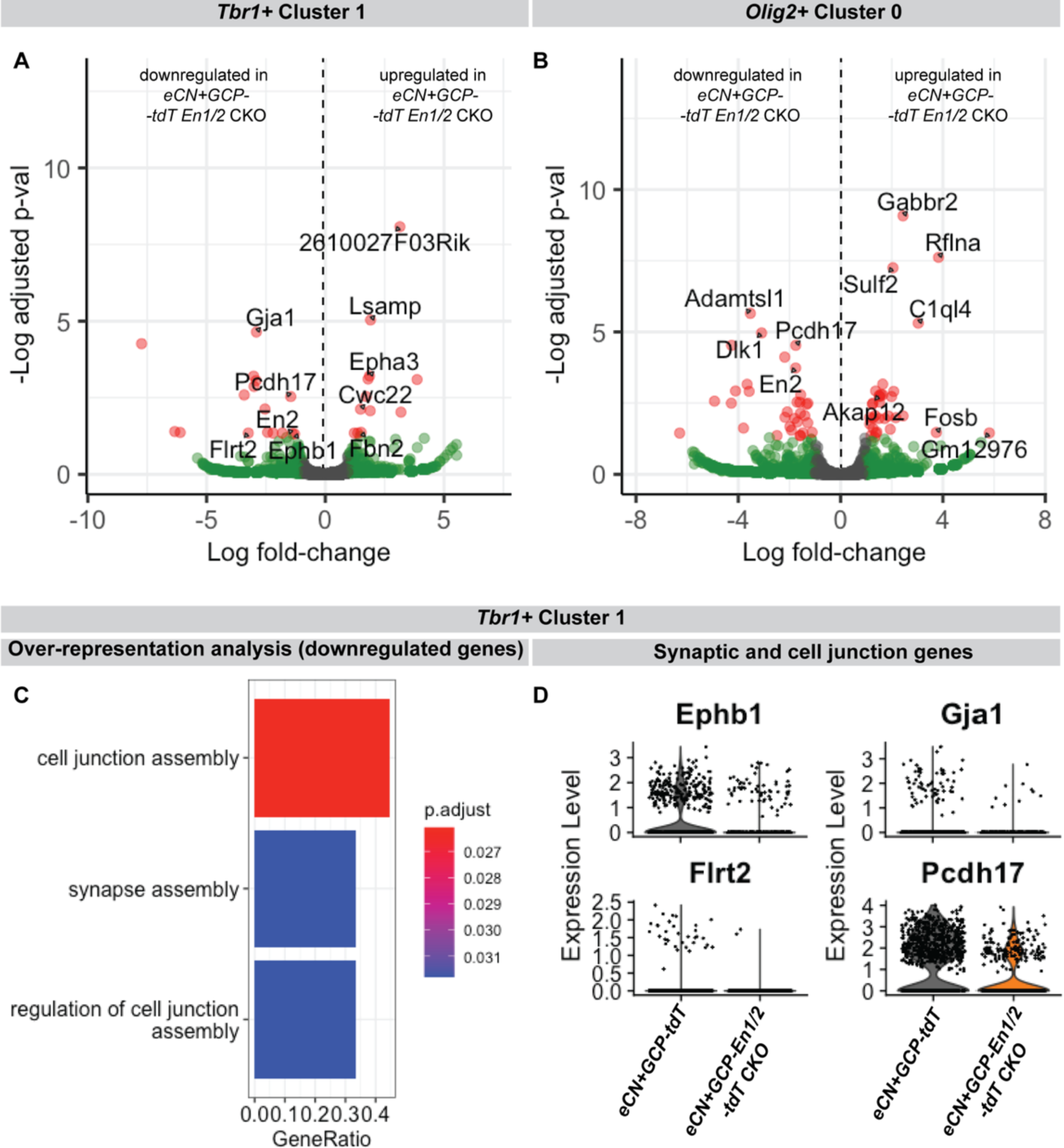
Differential gene expression analyses between *eCN+GCP-En1/2-tdT CKO* and *eCN+GCP-tdT* eCN samples. (A) Volcano plots depicting differentially expressed genes between *eCN+GCP-En1/2-tdT* CKO and *eCN+GCP-tdT* cells in *Tbr1+* cluster 1 of m+c analysis. Green dots indicate genes with log_2_ fold change > 2, red dots indicate genes significantly up- or downregulated with *P* ≤ 0.05 and log_2_ fold change > 2, black indicates non-significant genes. (B) Volcano plots depicting differentially expressed genes between *eCN+GCP-En1/2-tdT* CKO and *eCN+GCP-tdT* cells in *Olig2+* cluster 0 of m+c analysis. (C) Overrepresentation analysis of downregulated genes in *eCN+GCP-En1/2-tdT* CKO cells of *Tbr1+* cluster 1. (E) Violin plots depicting expression of the 4 differentially expressed genes associated with synaptic/cell junction function in *eCN+GCP-tdT* and *eCN+GCP-En1/2-tdT* CKO cells of *Tbr1+* cluster 1.

**Figure S8:**
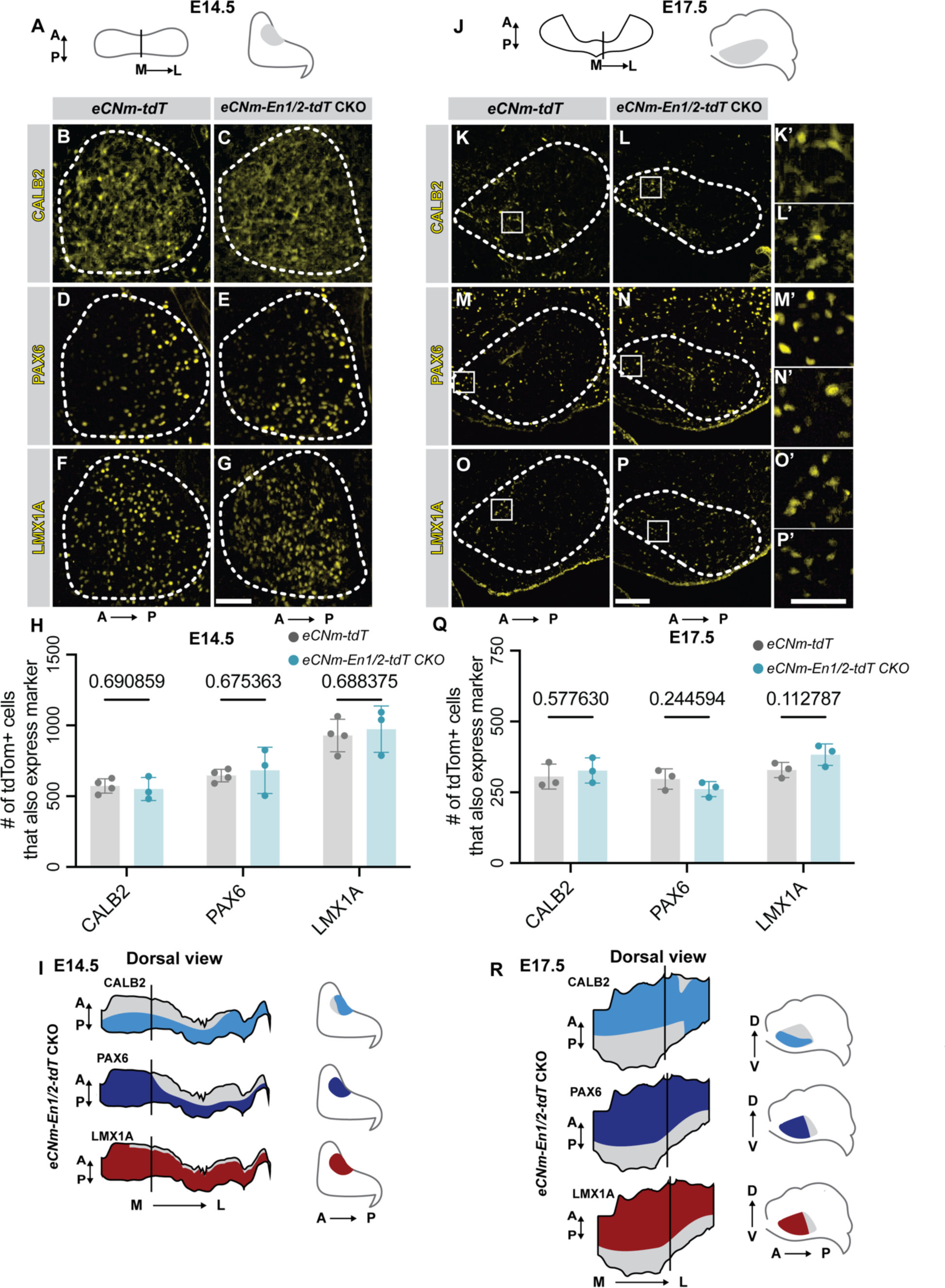
*En1/2* loss does not alter CALB2, PAX6 and LMX1A eCNm expression domains in *eCNm-En1/2-tdT* CKOs. (A) Schematic of E14.5 cerebellum, with lines indicating level at which (B - G) were taken. (B-G) Representative IF images of indicated proteins in eCNm of E14.5 mutants and controls. (H) Quantification of CALB2+, PAX6+, and LMX1A+ eCNm subpopulations in E14.5 mutant and control embryos (n = 4 *eCNm-tdT* and 3 *eCNm-En1/2-tdT* CKO animals, Student’s t-test). (I) Schematics of CALB2, PAX6 and LMX1A expression in eCNm of *eCNm-En1/2-tdT* CKOs projected onto dorsal view of 3D-reconstruction of the E14.5 eCNm. (J) Schematic of E17.5 cerebellum with lines indicating level at which (K-P) were taken. (K-P) Representative IF images of indicated proteins in eCNm of E17.5 mutants and controls. Rectangles indicate regions from which (K’-P’) were taken. (Q) Quantification of CALB2+, PAX6+ and LMX1A+ eCNm subpopulations in E17.5 mutant and control cerebella (n = 3 animals per genotype, Student’s t-test). (R) Schematics of CALB2, PAX6 and LMX1A expression in eCNm of *eCNm-En1/2-tdT* CKOs projected onto dorsal views of 3D-reconstruction of the E17.5 eCNm. Scale bars: 50 μm (B-I), 100 μm (K-P’).

**Figure S9:**
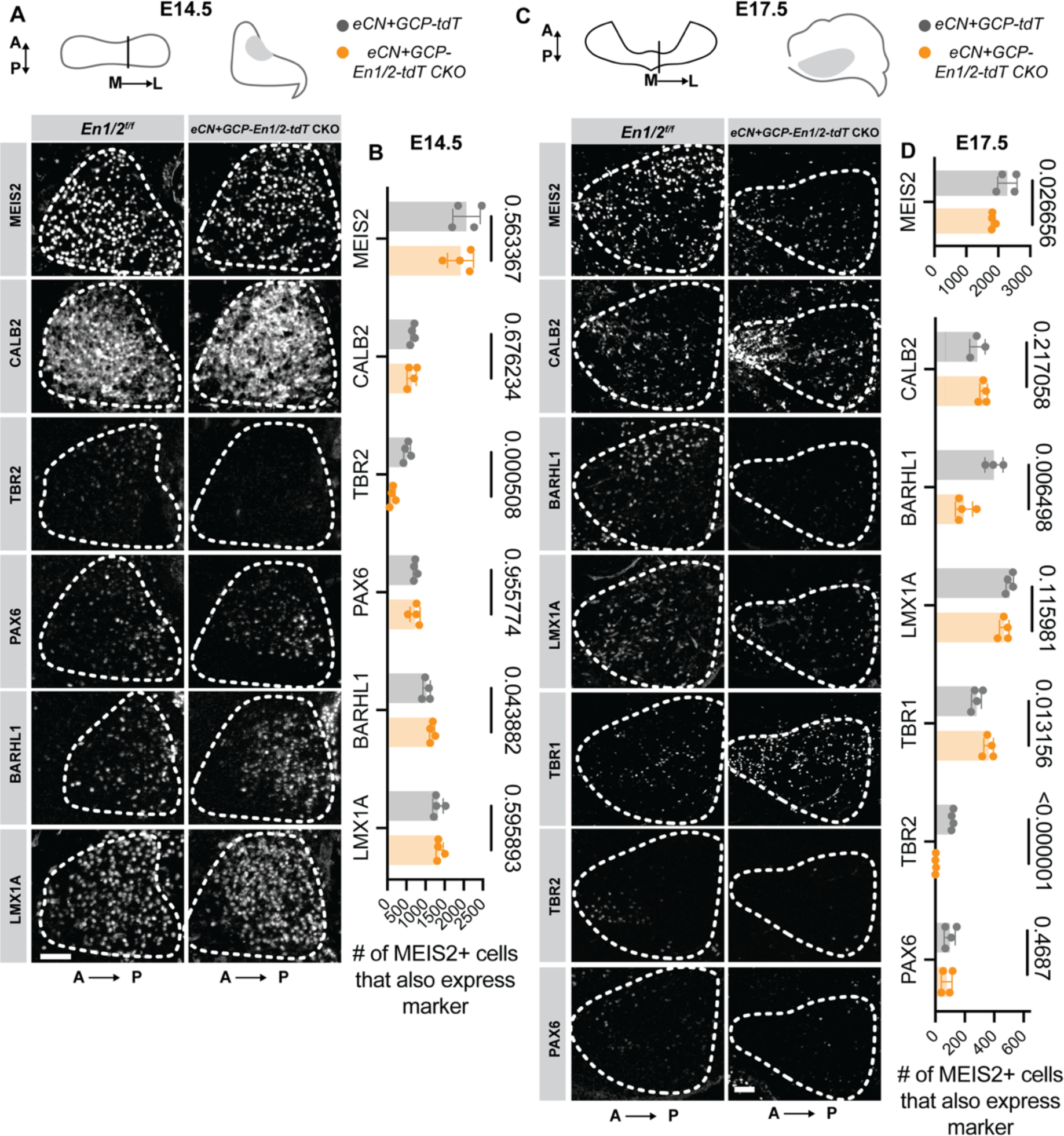
*En1/2* loss preferentially affects TBR2, BARHL1 and TBR1 expression domains in *eCN+GCP-En1/2-tdT* CKO mutants compared to controls. (A) Schematic of E14.5 cerebellum, with lines indicating level at which images below were taken of representative IF images of eCNm markers in E14.5 *eCN+GCP-En1/2-tdT* CKOs and littermate controls. (B) Quantification of MEIS2+ eCN that express eCNm markers in E14.5 *eCN+GCP-En1/2-tdT* CKO mutants and littermate controls. (n = 4 animals per genotype, Student’s t-test). (C) Schematic of E14.5 cerebellum, with lines indicating level at which images below were taken of representative IF images of eCNm markers in E17.5 mutants and controls. (B) Quantification of MEIS2+ eCN that express eCNm markers in E17.5 *eCN+GCP-En1/2-tdT* CKOs and littermate controls (n ≤ 3 animals per genotype, Student’s t-test). Scale bar: 50 μm (A), 100 μm (B).

**Figure S10:**
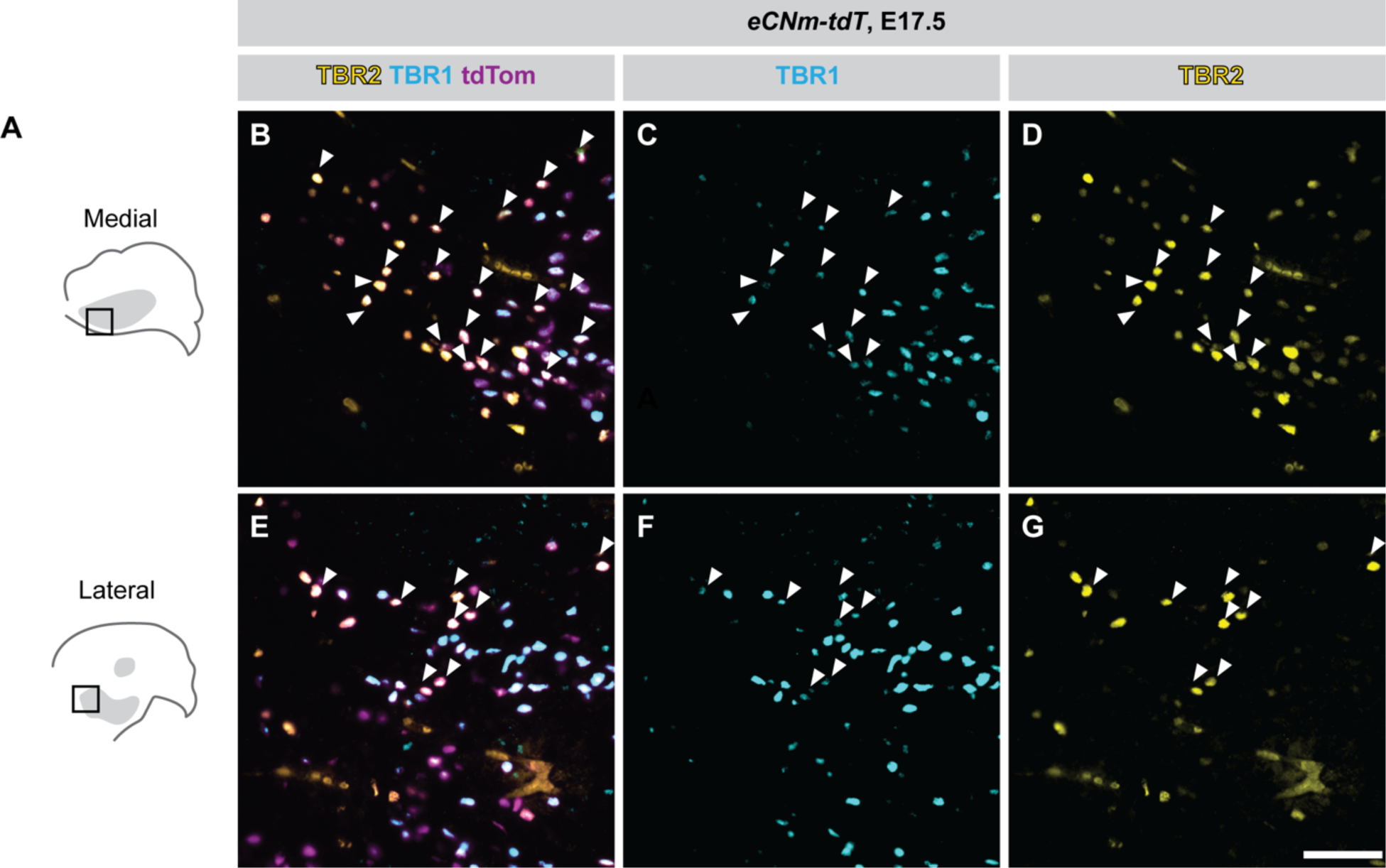
Anterior TBR2+ eCNm express TBR1 at E17.5. (A) Schematics of sagittal sections of the E17.5 cerebellum at medial (top) and lateral (bottom) levels of the eCNm with boxes indicating regions from which (B-D) and (E-G) were taken, respectively. Merge (B,E) or TBR1 (C,F) and TBR2 (D,G) single channel IF images of anterior eCNm in E17.5 *eCNm-tdT* embryos. White arrowheads indicate TBR1+ TBR2+ eCNm. Scale bar: 100 μm (B-G).

**Figure S11:**
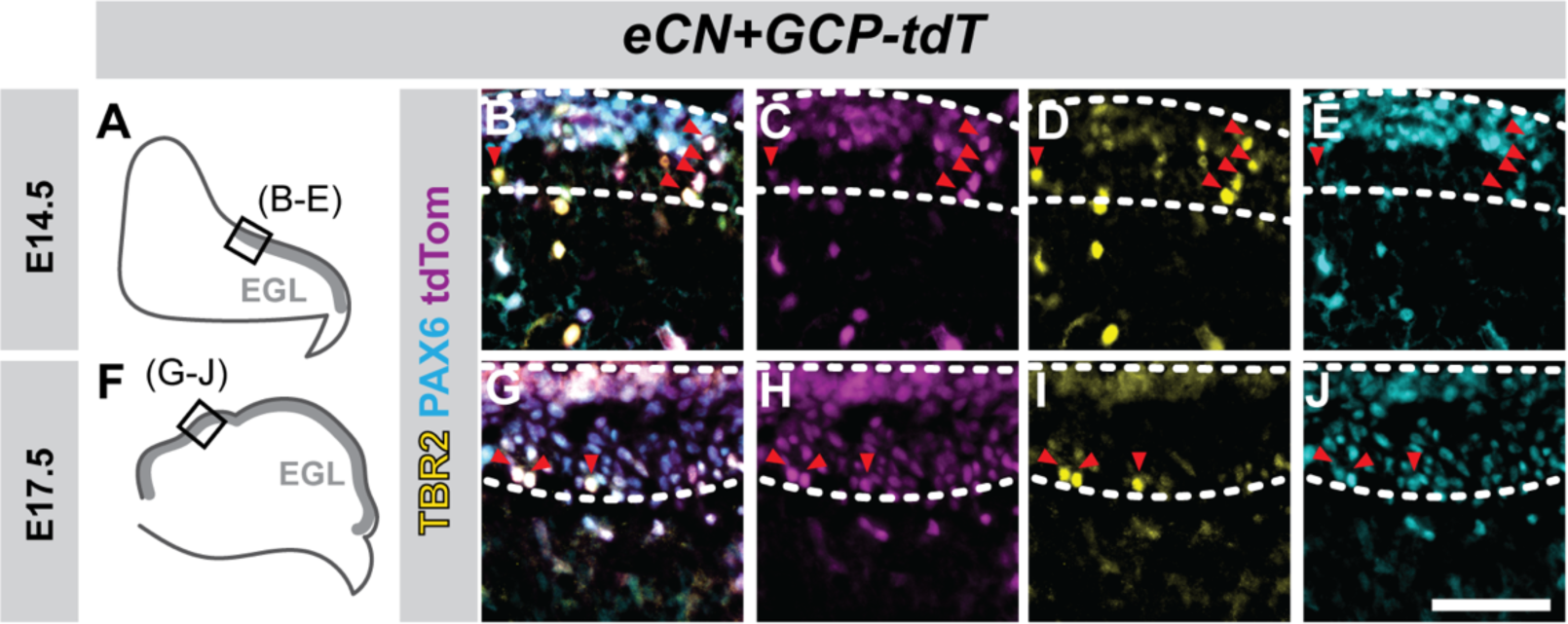
Expression of TBR2 in GCPs at E14.5 and E17.5. (A) Schematic of E14.5 sagittal cerebellum section with EGL depicted in grey. Box in (A) indicates region of EGL from which images in (B-E) were taken. Merge (B) or tdT (C), TBR2 (D) and PAX6 (E) single channel images of the EGL from E14.5 *eCN+GCP-tdT* cerebella with triple positive tdT+ TBR2+ PAX6+ GCPs indicated with red arrowheads. (F) Schematic of E17.5 sagittal cerebellum section with EGL depicted in light grey. Box in (F) indicates region of EGL from which (G-J) were taken. Merge (G) or tdT (H), TBR2 (I) and PAX6 (J) single channel images of the EGL from E17.5 *eCN+GCP-tdT* cerebella with triple positive tdT+ TBR2+ PAX6+ GCPs indicated with red arrowheads. Scale bar: 50 μm (B-J).

**Figure S12:**
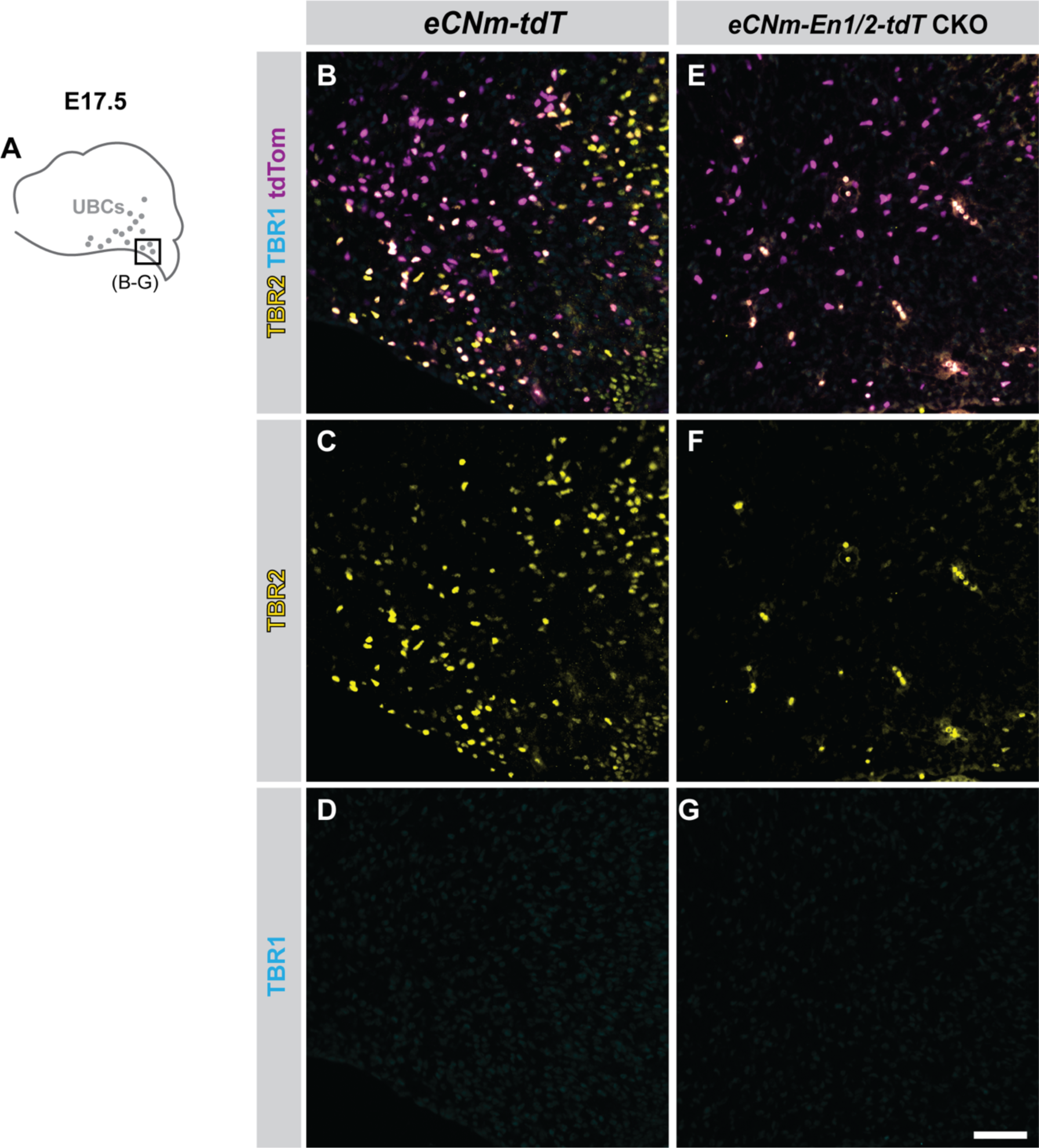
*En1/2* deficient UBCs do not express TBR2 at E17.5 and do not aberrantly upregulate TBR1. (A) Schematic of E17.5 sagittal cerebellum section with UBCs depicted by grey circles. Box in (A) indicates region from which images in (B-G) were taken. Merge (B, E) or TBR2 (C, F) and TBR1 (D, G) single channel images of UBCs in E17.5 *eCNm-tdT* and *eCNm-En1/2-tdT* CKO cerebella. Scale bar: 100 μm (B-G).

**Figure S13:**
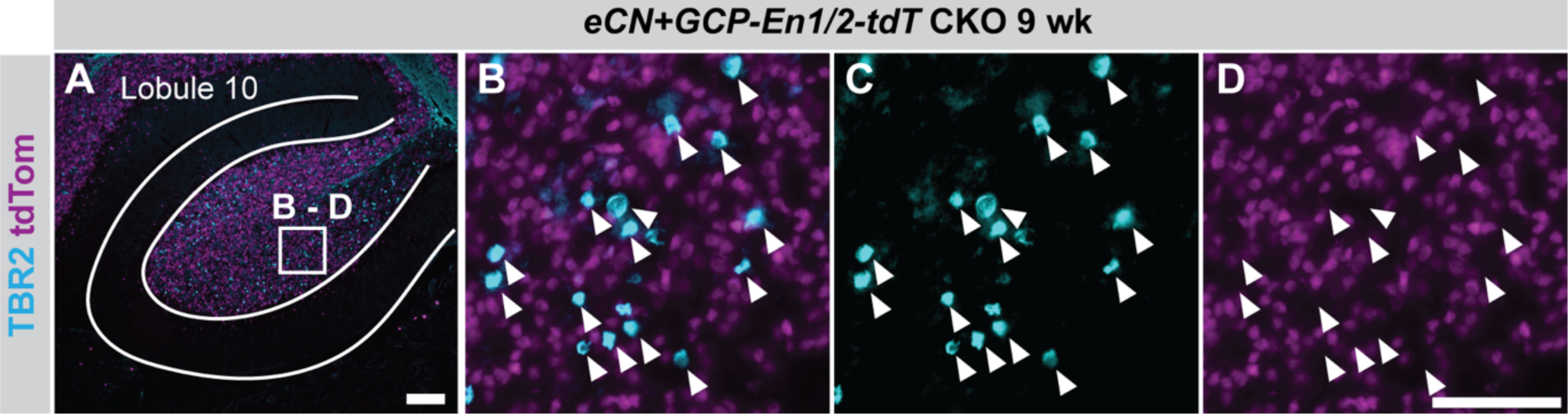
*eCN+GCP-Cre* does not recombine in UBCs, and UBCs in *eCN+GCP-En1/2-tdT* CKO animals are present. (A) Sagittal section of *eCN+GCP-En1/2-tdT CKO* cerebellum showing TBR2+ UBCs in lobule X. Merge (B) or single channel TBR2 (C) and tdT (D) high magnification images showing TBR2+ UBCs do not express tdT. Scale bars: 100 μm (A), 50 μm (B-D).

**Figure S14:**
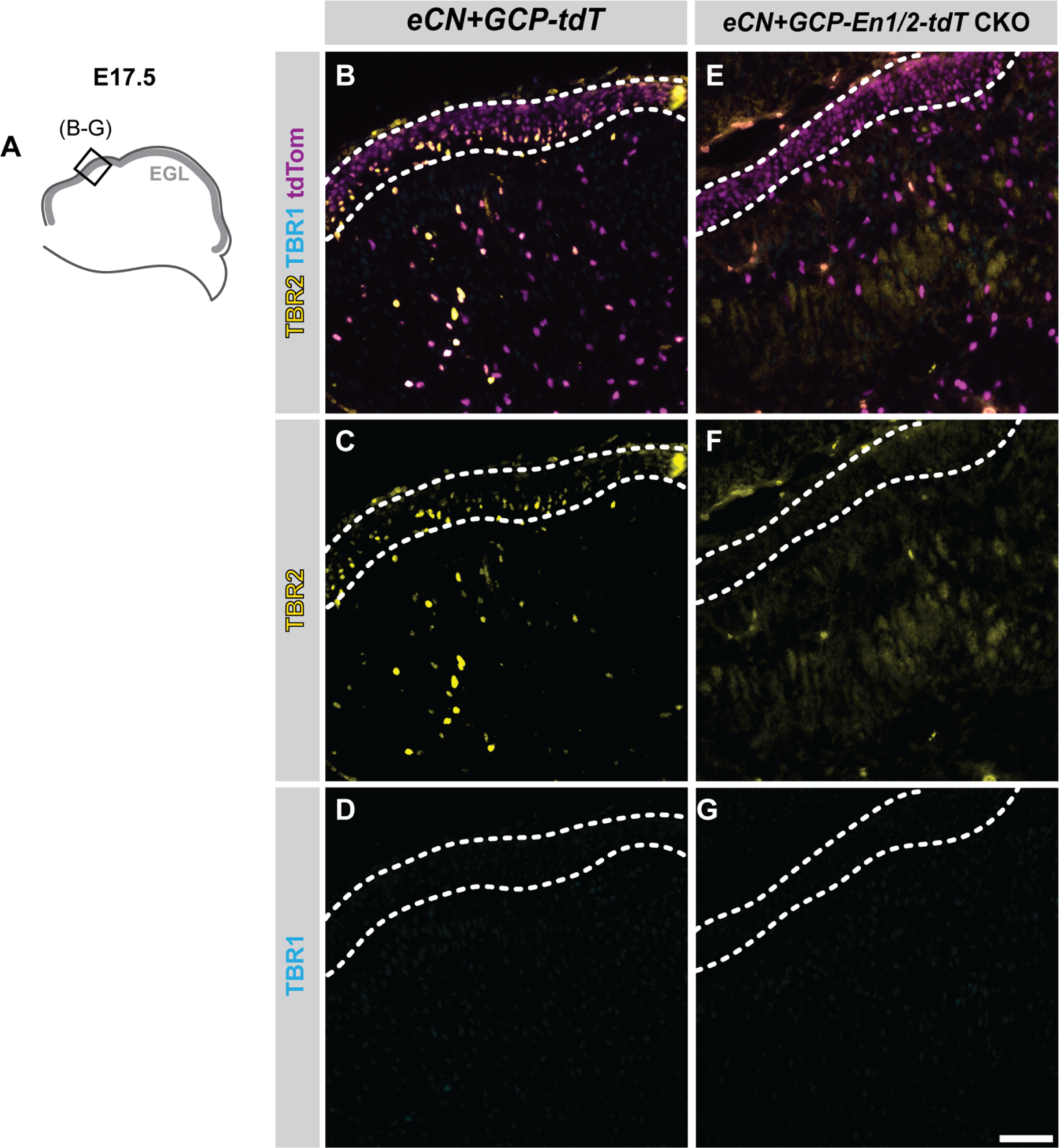
*En1/2* deficient GCPs down regulate TBR2 at E17.5 and do not aberrantly upregulate TBR1. (A) Schematic of E17.5 sagittal cerebellum section with EGL depicted by grey outline. Box in (A) indicates region from which images in (B-G) were taken. Merge (B, E) or TBR2 (C, F) and TBR1 (D, G) single channel images of GCPs in E17.5 *eCN+GCP-tdT* and *eCN+GCP-En1/2-tdT* CKO cerebella. Dashed outlines indicate EGL. Scale bar: 100 μm (B-G).

**Figure S15:**
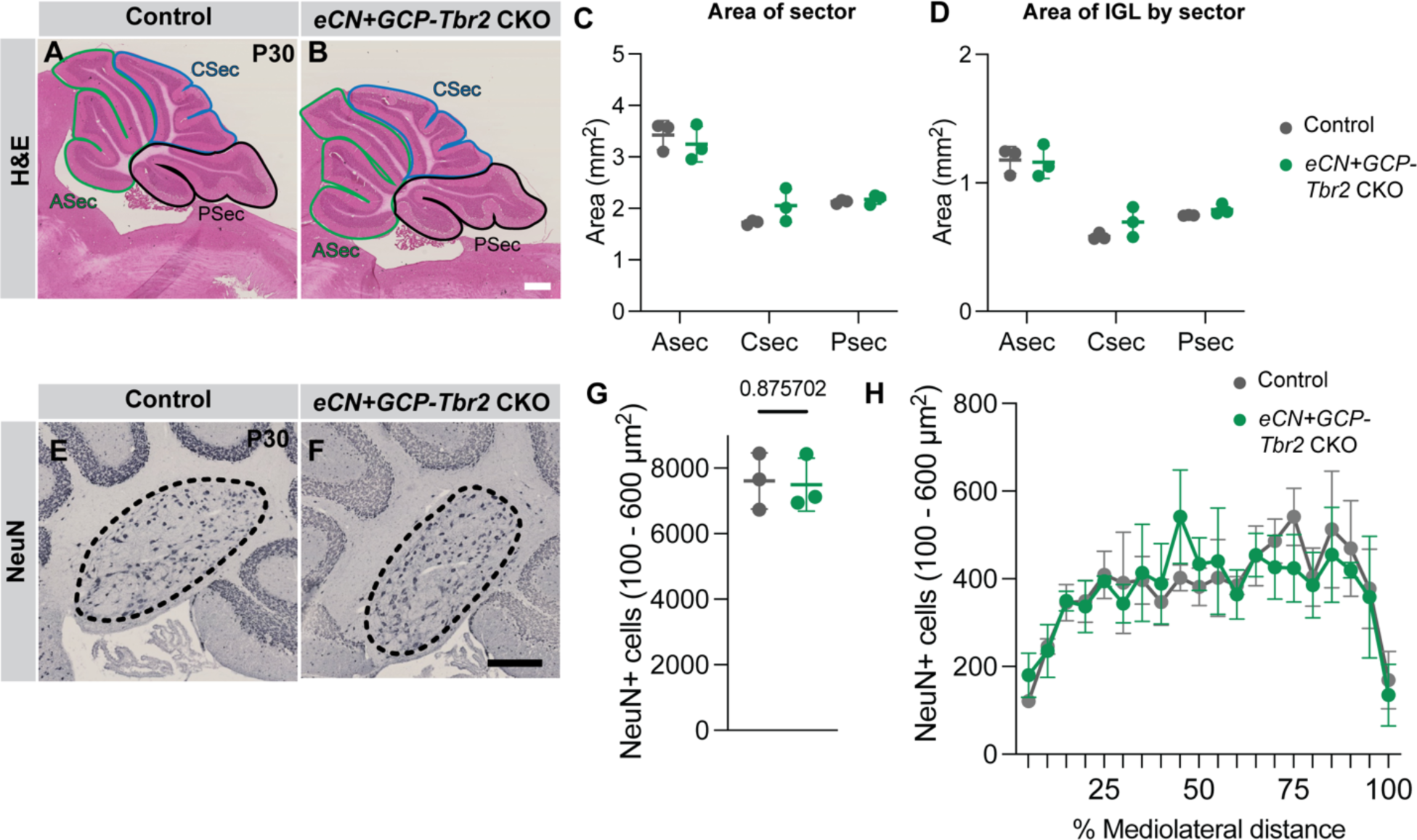
*Tbr2* conditional loss does not affect eCN and GCP survival or cerebellar growth and morphology. (A,B) Representative H & E images of sagittal sections of the vermis from P30 *eCN+GCP-Tbr2* CKO and control mice with sectors outlined. (C) Quantification of sector area showing no significant difference in *eCN+GCP-Tbr2* CKOs (n=3) compared to littermate controls (n = 2 *Tbr2^lox/+^* and n = 1 *Tbr2^lox/lox^*) (Two-way ANOVA, F_(1, 12)_ = 0.3708). (D) Quantification of IGL area by sector showing no significant difference in *eCN+GCP-Tbr2* CKOs (F_(1, 12)_ = 1.394). (E-F) Representative NeuN images of the eCNm at P30 in *eCN+GCP-Tbr2* CKO and littermate control. (G) Quantification of NeuN+ cells that are 100–600 um^2^ (eCN size range) showing no reduction in *eCN+GCP-Tbr2* CKOs compared to littermate controls. (H) Mediolateral distribution of 100–600 um^2^ NeuN+ cells showing no change in distribution of eCN across mediolateral axis in *eCN+GCP-Tbr2* CKOs compared to littermate controls. Scale bars: 500 um (A,B), 250 um(E,F). ASec: anterior sector; CSec: central sector; PSec: posterior sector.

## Supplemental Table titles (see excel file with 13 tabs)

**Table S1**. Significantly enriched genes (adjusted p value ≤ 0.05) for each tdt+ cluster from *eCN+GCP-tdT* clustering. Related to Figure 2 and Figure S2.

**Table S2**: Significantly enriched genes (adjusted p value ≤ 0.05) for each *Meis2-*enriched cluster from *eCN+GCP-tdT* subclustering. Related to Figure 2 and Figure S2.

**Table S3**: Significantly enriched genes (adjusted p value ≤ 0.05) for each eCN cluster from *eCN+GCP-tdT* subclustering. Related to Figure 2 and Figure S2.

**Table S4**: Significantly enriched genes (adjusted p value ≤ 0.05) for each tdT+ cluster from *eCN+GCP-En1/2-tdT* CKO clustering. Related to Figure S5.

**Table S5:** Significantly enriched genes (adjusted p value ≤ 0.05) for each *Meis2-* enriched cluster from *eCN+GCP-En1/2-tdT* CKO subclustering. Related to Figure S5.

**Table S6:** Significantly enriched genes (adjusted p value ≤ 0.05) for each eCN cluster from *eCN+GCP-En1/2-tdT* CKO subclustering. Related to Figure S5.

**Table S7**: Significantly enriched genes (adjusted p value ≤ 0.05) for each tdT+ cluster from *eCN+GCP-En1/2-tdT* CKO and *eCN+GCP-tdT* combined clustering. Related to Figure 5 and Figure S6.

**Table S8:** Significantly enriched genes (adjusted p value ≤ 0.05) for each *Meis2-* enriched cluster from *eCN+GCP-En1/2-tdT* CKO and *eCN+GCP-tdT* combined subclustering. Related to Figure 5 and Figure S6.

**Table S9:** Significantly enriched genes (adjusted p value ≤ 0.05) for each eCN cluster from *eCN+GCP-En1/2-tdT* CKO and *eCN+GCP-tdT* combined subclustering. Related to Figure 5 and Figure S6.

**Table S10:** Differentially expressed genes (adjusted p value ≤ 0.05) between eCN of *eCN+GCP-En1/2-tdT* CKO mutants and *eCN+GCP-tdT* controls. Related to Figure S6.

**Table S11:** Significantly enriched GO terms (adjusted p value ≤ 0.05) from over-representation analysis (downregulated genes in *eCN+GCP-En1/2-tdT* CKO mutants compared to controls) in Tbr1+ eCNm. Related to Figure S6.

**Table S12:** Differentially expressed genes (adjusted p value ≤ 0.05) between GCPs (clustered by cell cycle phase) of eCN*+GCP-En1/2-tdT CKO* mutants and *eCN+GCP-tdT* controls. Related to Figure 8.

**Table S13:**
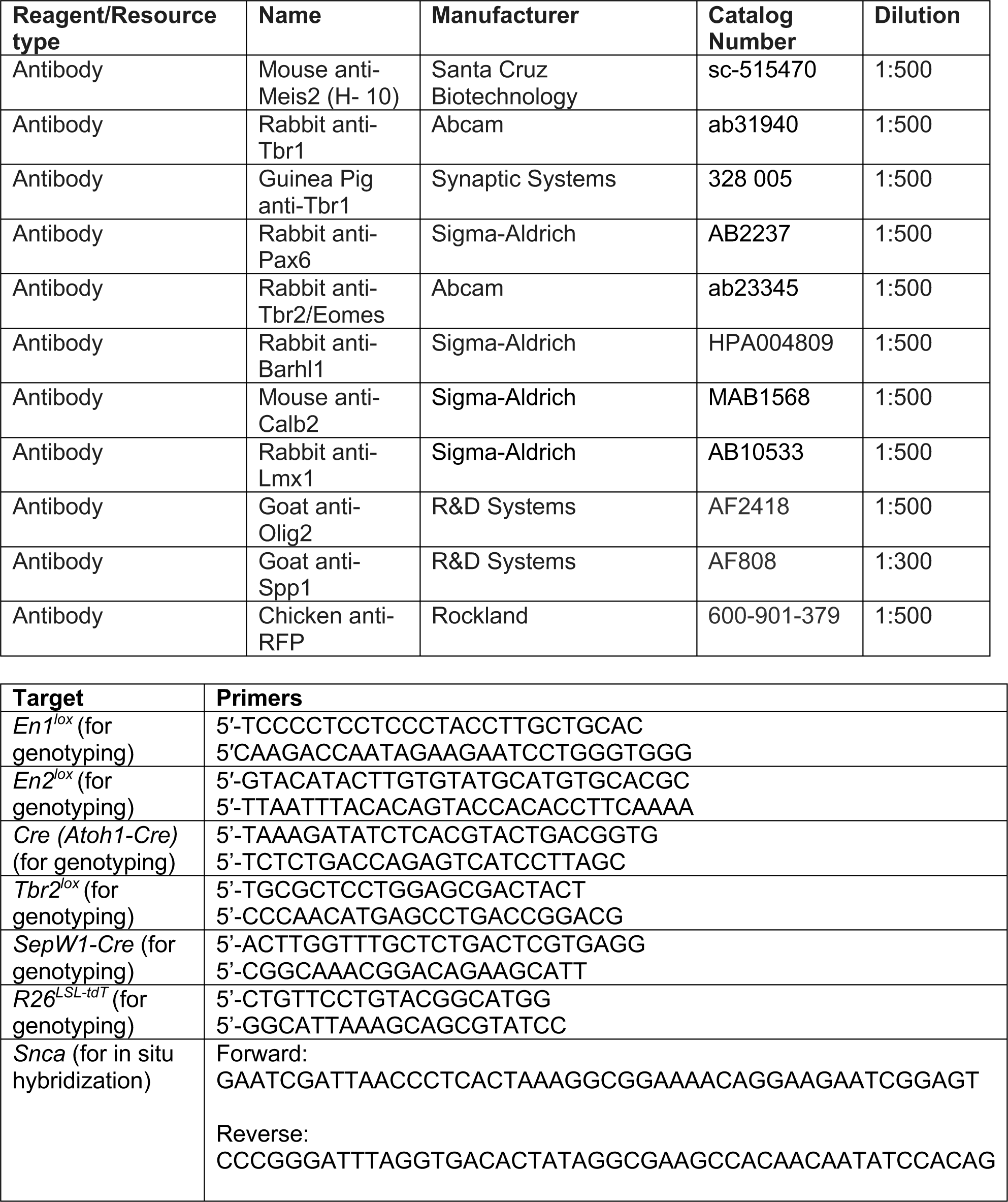
Significantly enriched GO terms (adjusted p value ≤ 0.05) from over-representation analysis (downregulated genes in *eCN+GCP-En1/2-tdT* CKO mutants compared to controls) in G0/G1 phase GCPs. Related to Figure 8.

**Table S14: List of antibodies and primers used in this study.**

## Notes

### Competing Interest Statement

The authors have declared no competing interest.

